# MM-ComBat and MM-CovBat: Multivariate Frameworks for Joint Harmonization of Multi-Metric Neuroimaging Data

**DOI:** 10.64898/2026.02.05.704069

**Authors:** Zheng Ren, Patrick Sadil, Martin A. Lindquist

**Affiliations:** Department of Biostatistics, Johns Hopkins Bloomberg School of Public Health, 615 N. Wolfe Street, Baltimore, Maryland 21205, USA

**Keywords:** neuroimaging, harmonization, ComBat, multivariate

## Abstract

Aggregating neuroimaging data across sites and studies is increasingly common, yet site- and scanner-related batch effects can obscure meaningful biological variation and introduce spurious associations. Although ComBat and its extensions are widely used, they are primarily designed for single-metric harmonization. In practice, neuroimaging studies often involve multiple biologically coupled metrics (e.g., cortical thickness, surface area, and gray-matter volume) measured across multiple features (e.g., regional values), with shared covariance structure both within and across metrics. Applying single-metric ComBat independently to each metric ignores these cross-metric dependencies. Using data from the NIH Acute to Chronic Pain Signatures (A2CPS) program, we show that batch effects occur not only in means and variances but also in covariance across cortical regions and metrics—relationships that single-metric ComBat does not fully remove. We propose MM-ComBat, a multivariate extension of ComBat that jointly harmonizes multiple metrics by borrowing strength across them, better capturing cross-metric dependence while also improving covariance estimation across features. Because joint harmonization whitens residual covariance toward a standardized baseline, it risks distorting biologically meaningful cross-metric structure when batch effects are moderate. We therefore introduce two complementary formulations: a baseline formulation suited to batch-dominated settings, and a target-covariance formulation that remaps adjusted covariances toward an estimated shared biological structure rather than fully whitening them. Both empirical Bayes (EB) and Bayesian Markov Chain Monte Carlo (MCMC) implementations of MM-ComBat effectively reduce batch effects. In our experiments, EB is more robust to measurement error, whereas MCMC more accurately recovers cross-metric correlations when priors are well specified. Recognizing that batch effects can also affect feature-level covariance, CovBat was recently introduced as an extension of ComBat that harmonizes both first- and second-order moments across sites. We extend CovBat to the multivariate framework as MM-CovBat, which performs a second-stage latent-space harmonization to directly address covariance-related batch effects across features and metrics. Simulations confirm that MM-ComBat improves correlation recovery and better preserves biological effects in the mean structure relative to single-metric ComBat, particularly for moderate-to-strong effects, and that MM-CovBat further improves separation of true biological variation from batch effects when independence assumptions are violated. Together, these methods provide a flexible and unified framework for harmonizing complex, multi-metric neuroimaging data in large-scale, multi-site studies.

## 1 Introduction

Large-scale neuroimaging research now routinely requires pooling data across sites and scanners to achieve adequate sample sizes and improve generalizability. There is also growing interest in integrating multi-domain data to enable comprehensive biomarker discovery. One initiative exemplifying this trend is the National Institutes of Health (NIH)-funded *Acute to Chronic Pain Signatures (A2CPS)* program, which aims to identify and validate biomarkers of pain chronification by jointly analyzing neuroimaging, genomic, and behavioral measures, among others (Sluka et al. 2023). Such large-scale, multi-site, and multi-modal efforts highlight both the promise of integrative analyses and the challenges posed by technical and site-specific variability. Imaging data are particularly susceptible, because heterogeneous acquisition protocols and scanner parameters can introduce site- and scanner-related differences, commonly referred to as batch effects. These effects are not biologically meaningful, yet they can confound associations of interest, reduce reproducibility, and limit interpretability. Effective harmonization methods are therefore essential to remove non-biological variability while preserving biologically meaningful signal.

Among the harmonization approaches that have been developed, ComBat has become one of the most widely adopted. Originally proposed for genomic expression arrays (Johnson, Li, and Rabinovic 2006), it has since been successfully applied across diverse neuroimaging modalities including diffusion tensor imaging, cortical thickness, functional connectivity, and radiomic features derived from positron emission tomography (Fortin, Parker, et al. 2017; Fortin, Cullen, et al. 2018; Yu et al. 2018; Orlhac et al. 2018). By pooling information across features to estimate prior distributions on batch effects (an empirical Bayes strategy), ComBat achieves stable, shrinkage-regularized estimates of site-related shifts in mean and variance even when individual feature-level estimates are noisy. This regularization is what allows ComBat to remove technical variability while preserving biological associations, and it is what makes the framework naturally extensible. Several extensions have been proposed to accommodate more complex data structures, including ComBat-GAM (Pomponio et al. 2020) for nonlinear covariate effects, LongComBat (Beer et al. 2020) for longitudinal data, and CovBat (Chen, Beer, et al. 2021) for harmonizing feature covariance across batches to enhance downstream machine learning analyses. More recently, a fully Bayesian version of ComBat has been introduced (Reynolds et al. 2023), which may better preserve biological variation while achieving more accurate harmonization.

Before describing our approach, we clarify two terms used throughout. A *feature* denotes a spatial or anatomical unit on which measurements are recorded (e.g., a cortical region), and a *metric* denotes a type of imaging-derived measurement recorded for each feature. Despite the advances described above, traditional ComBat and its variants are typically applied to one metric at a time. However, neuroimaging analyses often involve multiple biologically related metrics derived either from the same imaging modality (e.g., cortical thickness, surface area, and gray-matter volume from T1-weighted MRI) or across modalities (e.g., T1/T2 ratios and diffusion-derived measures). Measurements across these metrics often exhibit structured covariance both within and across metrics, reflecting shared underlying biological processes. For example, cortical thickness and surface area follow a dynamic relationship across the adult lifespan (Storsve et al. 2014), and morphometric similarity network analyses suggest macroscale co-alteration patterns across regions (Lu et al. 2024). More recently, Sadikov et al. (2025) demonstrated that diffusion MRI yields dozens of reliably measurable cortical microstructural metrics that exhibit substantial shared variance, which can be summarized by a smaller number of latent factors. Importantly, because these metrics are derived from the same images acquired on the same scanners, batch effects may also share common structure across metrics. This creates an opportunity to borrow information across them to improve estimation and removal. Applying single-metric ComBat independently to each metric ignores these cross-metric dependencies, leaving residual batch effects in covariance structure and potentially attenuating true biological variation.

To address this limitation, we propose MM-ComBat, a multivariate extension of the ComBat framework that jointly harmonizes multiple metrics by borrowing strength across correlated features and metrics. Unlike single-metric approaches, MM-ComBat explicitly models shared covariance structure, enabling more effective removal of batch effects in both means and covariances while preserving biologically meaningful relationships in the mean structure. However, because this joint modeling whitens residual covariance toward a standardized baseline, it risks distorting biologically meaningful cross-metric covariance when batch effects are moderate and biological covariance is strong. We therefore also introduce a target-covariance mapping strategy, which remaps adjusted covariances toward an estimated shared target rather than fully whitening them. Together, the baseline and target-covariance formulations offer complementary strategies. The former is preferable when batch-related covariance is dominant, the latter when preserving biological covariance structure is the priority. This framework is applicable both to studies investigating joint associations across metrics and to metric-specific analyses, where borrowing information across correlated metrics can stabilize batch effect estimation.

MM-ComBat, however, assumes that batch effects are independent across features and therefore does not fully correct structured batch effects in the cross-feature covariance space. To address this, we extend CovBat (which harmonizes both first- and second-order moments across sites) to the multivariate setting, introducing MM-CovBat. Where MM-ComBat adjusts batch effects within each feature’s cross-metric covariance, MM-CovBat additionally targets batch effects that persist in the cross-feature covariance structure, via a second-stage harmonization step in a shared latent space. This makes MM-CovBat particularly effective when the independence assumption is violated (i.e., when batch effects are structured across features rather than acting on each feature separately).

We demonstrate the utility of these methods using data from the A2CPS project. MM-ComBat more effectively reduces batch effects than single-metric ComBat both within and across metrics, with the largest gains for noisy metrics where borrowing information across correlated metrics stabilizes batch-effect estimation. MM-CovBat further improves separation of biological variation from batch effects, achieving the best overall performance, particularly when batch effects are structured across features. Between the two covariance formulations, target-covariance mapping is preferable when biological covariance is strong and batch effects are moderate, as the baseline formulation risks whitening meaningful biological structure along with batch-related variation. However, this approach may also reintroduce batch effects if the estimated target-covariance retains residual batch-related structure, indicating a trade-off between batch removal and preservation of biological covariance patterns. When batch-related covariance dominates, the baseline formulation performs better. Simulation studies corroborate these findings and confirm that the methods better preserve biological signals in the mean structure and more accurately recover true correlation structure relative to single-metric approaches, and further confirm that target-covariance mapping is preferable when strong biological structure remains in cross-metric covariance and batch effects are moderate or absent, while the baseline approach performs better when batch-related covariance structure is strong and biological covariance patterns are weak.

The remainder of the paper is organized as follows. Section 2 describes the A2CPS dataset, reviews existing ComBat-based methods, derives MM-ComBat and MM-CovBat, and illustrates the simulation study designs. Section 3 defines the harmonization evaluation criteria and Sections 4 and 5 present the empirical and simulation results. Finally, Section 6 discusses limitations and directions for future work.

## 2 Method

### 2.1 A2CPS dataset

The data analyzed in this study consist of baseline scans (release 1.0.0) from the Acute to Chronic Pain Signatures (A2CPS) program, funded by the National Institutes of Health (NIH) Common Fund. A2CPS aims to identify biomarkers and biosignatures across multiple data types that, individually or jointly, predict susceptibility or resilience to developing chronic pain after an acute pain event (Sluka et al. 2023). Informed consent was obtained from all participants. Detailed inclusion criteria and imaging acquisition protocols are described in Berardi et al. (2022).

A2CPS imaging data are collected across multiple sites using scanners from General Electric (GE), Siemens, and Philips, with protocols adapted from the Adolescent Brain Cognitive Development (ABCD) study; see Sadil et al. (2024) for more details. In this work, we focus on T1-weighted structural MRI data that provide multiple imaging-derived metrics across cortical regions. Each A2CPS T1-weighted scan received a quality rating on a three-point scale (“green”, “yellow”, “red”), where “yellow” and “red” indicate potential quality concerns. Accordingly, only scans rated “green” were included in this study.

Preprocessing was performed using *fMRIPrep* (Esteban et al. 2018). The T1-weighted (T1w) images were corrected for intensity non-uniformity (INU) using `N4BiasFieldCorrection` (Tustison et al. 2010) and served as the anatomical reference throughout the workflow. Skull stripping was performed using a Nipype implementation of the ANTs brain extraction workflow (`antsBrainExtraction.sh`), with the OASIS30ANTs template as the target (Avants et al. 2011). Cortical surface reconstruction was carried out using FreeSurfer (`recon-all`; Fischl 2012).

Harmonization performance of existing methods may decline as the number of features increases, likely due to the increasingly complex covariance structures across features and potential confounding with biological variables. Prior studies have shown that, for covariance-based harmonization methods such as CovBat, site effects may become more detectable after harmonization in settings with relatively small sample sizes and high-dimensional feature spaces, potentially due to poor covariance estimation (Chen, Beer, et al. 2021). To rigorously evaluate harmonization performance across methods, we therefore consider a deliberately challenging scenario. Specifically, we focus on the Destrieux cortical parcellation, which includes 78 regions of interest (ROIs), each characterized by eight different cortical metrics: surface area (SurfArea), gray-matter volume (GrayVol), mean thickness (ThickAvg), thickness standard deviation (ThickStd), mean curvature (MeanCurv), Gaussian curvature (GausCurv), folding index (FoldInd), and curvature index (CurvInd). These metrics include quantities with known mathematical and biological relationships, as well as differing statistical scales, thereby providing a stringent test of harmonization methods in high-dimensional and correlated settings.

The analytic sample included 473 participants (34% male; mean age = 63 years; mean self-reported pre-surgical pain duration = 53 months) from six sites and three scanner manufacturers: Wayne State (WS; Siemens, *n* = 29), University of Illinois Chicago (UI; GE, *n* = 130), University of Michigan (UM; GE, *n* = 97), Spectrum Health (SH; Siemens, *n* = 22), University of Chicago (UC; Philips, *n* = 82) and Endeavor Health (NS; Siemens, *n* = 113). Site and scanner differences represent a major source of potential batch effects. Effective harmonization is essential to mitigate these effects while preserving biologically relevant variation, thereby improving the reliability of downstream analyses and predictive modeling.

### 2.2 Review of ComBat-based methods

We first review commonly used ComBat-based methods for harmonizing neuroimaging data. The original ComBat (Johnson, Li, and Rabinovic 2006) and its extensions aim to remove batch effects in feature-wise mean and variance within an empirical Bayes (EB) framework. Here, fixed covariate effects are estimated and preserved via regression, while batch-specific location and scale parameters are estimated via EB posterior means, with prior hyperparameters obtained using method-of-moments (MoM) estimation across features.

The methods vary primarily in how they model the underlying biological signal and data structure. *ComBat* assumes cross-sectional data with approximately linear covariate effects and harmonizes each feature independently. *LongComBat* extends this framework to longitudinal data by incorporating subject-specific random effects, *ComBat–GAM* uses smooth functions to capture nonlinear covariate effects, and *CovBat* further adjusts the covariance structure across features to reduce batch-related differences in multivariate correlations.

#### Original ComBat

The original *ComBat* is designed for cross-sectional data with approximately linear covariate effects. Let *Y*_*ijv*_ denote the value of feature *v* for subject *j* in batch *i*, and let ***X***_*ij*_ be a vector of covariates. *ComBat* models each feature as

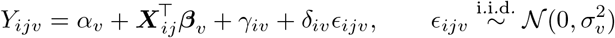

where *α*_*v*_ is a feature-specific intercept, ***β***_*v*_ are fixed-effect coefficients, and *γ*_*iv*_ and *δ*_*iv*_ represent batch-specific location (mean) and scale (variance) effects. For each feature *v*, least-squares estimates 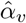 and 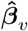 are obtained by regression. Empirical Bayes priors are then placed on the batch effects across features, with hyperparameters estimated by MoM. Posterior means 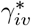 and 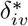 are used to adjust the data while preserving the fixed effects, yielding the harmonized values:

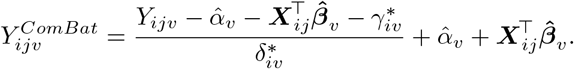

#### LongComBat

*LongComBat* extends the original ComBat framework to longitudinal data, where participants have repeated measurements over time. For feature *v* of subject *j* in batch *i* at time *t*, the model is

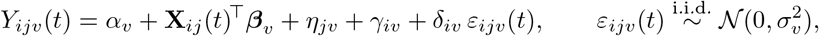

where *η*_*jv*_ is a subject-specific random effect, *γ*_*iv*_ and *δ*_*iv*_ are batch-specific location and scale effects, and **X**_*ij*_(*t*) contains biological covariates to be preserved. Fixed and random effects are first estimated using restricted maximum likelihood (REML). Similar to *ComBat*, batch effects are then estimated in an EB framework, with hyperparameters obtained via MoM, and posterior means of the batch effects subsequently used for adjustment. The harmonized values are

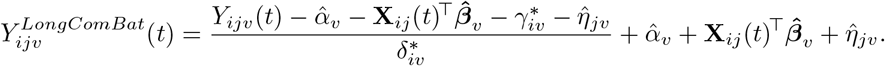

This formulation enables harmonization of repeated measures while accounting for within-subject correlations and between-site variability.

#### ComBat-GAM

For cross-sectional data with nonlinear covariate effects, *ComBat–GAM* replaces the linear term with a smooth function:

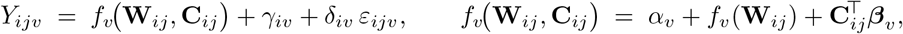

where *f*_*v*_(·) is a spline-based smooth function of the nonlinear covariates **W**_*ij*_ for feature *v*, and **C**_*ij*_ contains covariates modeled linearly. The smooth function 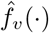 is estimated via generalized additive models (GAM), after which the standard *ComBat* EB pipeline provides posterior means 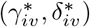 for the batch effects. The harmonized values are

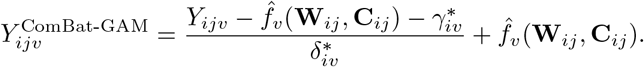

By fitting nonlinear biological effects before batch correction, ComBat-GAM avoids conflating smooth covariate trends with site-related shifts in mean and variance.

#### CovBat

The three methods above primarily focus on batch effects in feature-wise means and scales. However, they do not correct batch effects in cross-feature covariance, which can bias association estimates and degrade downstream machine learning performance. *CovBat* extends the *ComBat* framework to harmonize covariance through a two-stage procedure.

##### Stage 1 (Feature-wise ComBat)

Standard ComBat is first applied to remove batch-specific mean and variance effects from each feature’s marginal distributions, providing residuals

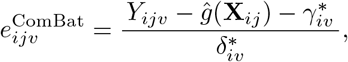

so that, for each batch *i*, the residual vector 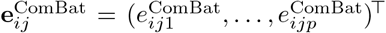 has mean zero and batch-specific covariance **Σ**_*i*_. Here, *p* is the total number of features, and *ĝ*(·) denotes the estimated contribution of the preserved biological effects.

##### Stage 2 (PCA-domain covariance harmonization)

Principal component analysis (PCA) is performed on the pooled residual covariance matrix to obtain eigenvectors 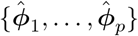 and principal component (PC) scores 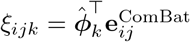. The first *K* score coordinates are harmonized across batches to produce 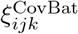, and the harmonized residuals are reconstructed as:

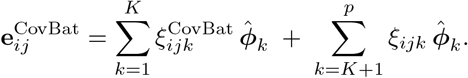

Here *K* controls the fraction of total variance whose covariance structure is harmonized. Finally, the preserved fixed effects are added back to obtain CovBat-adjusted observations:

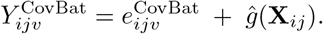

This two-stage approach harmonizes both marginal distributions and the cross-feature covariance across batches, improving consistency for downstream multivariate analyses.

### 2.3 Multi-metric ComBat

The four ComBat-based methods described above have demonstrated promising harmonization performance in single-metric settings, which we collectively refer to as the single-metric ComBat (SM-ComBat) framework. However, many neuroimaging studies analyze multiple correlated metrics, such as cortical thickness, surface area, and curvature, which may share both biologically meaningful covariance and batch-related covariance across metrics and regions. Applying SM-ComBat independently to each metric ignores this cross-metric dependence structure and may therefore inadequately address covariance-related batch effects, potentially leaving residual batch effects in cross-metric covariance or distorting shared biological patterns across metrics.

To address this, we propose two joint covariance harmonization approaches, which we collectively refer to as multi-metric ComBat (MM-ComBat). The first assumes that biological signal is captured primarily in the mean structure, so that the residuals mainly reflect shared batch structure and noise. Under this assumption, residual covariance is mapped toward a standardized reference to remove batch-related structure. We refer to this version as MM-ComBat (baseline covariance). The second assumes that biological signal persists in both mean and covariance across metrics, so that the residual covariance contains a mixture of biological structure, batch-related variation, and noise. Under this formulation, batch-specific covariance is mapped toward a shared target estimated from the data, with the goal of preserving common biological structure. We refer to this version as MM-ComBat (target covariance).

Broadly speaking, the baseline variant prioritizes aggressive removal of cross-metric batch structure, while the target variant trades some removal for better preservation of shared biological covariance. Because the baseline variant maps all residual covariance toward a standardized reference, it risks removing biologically meaningful cross-metric structure alongside batch-related variation when biological covariance is strong and batch effects are moderate — a risk we refer to throughout as the *whitening concern*. Accordingly, we first present MM-ComBat (baseline covariance) and then describe the covariance reintroduction step as an additional component of MM-ComBat (target covariance).

Building on the principles of existing ComBat methods, MM-ComBat assumes that, after removing fixed effects, the residuals follow a multivariate normal distribution and are independent across observations within each batch. This extension allows MM-ComBat to jointly model multiple correlated metrics rather than treating them separately, thereby enabling batch adjustment of both marginal distributions and cross-metric covariance structure.

#### 2.3.1 Multivariate Framework Setup

For simplicity, we introduce the MM-ComBat framework under a linear model. For each batch *i*, feature *v*, and metric *m*, we assume

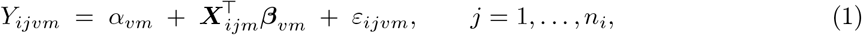

where *α*_*vm*_ is a metric-specific intercept, ***X***_*ijm*_ is the covariate vector, ***β***_*vm*_ are the corresponding fixed-effect coefficients, *ε*_*ijvm*_ is the residual error term, and *n*_*i*_ is the number of subjects in batch *i*. We first estimate *α*_*vm*_ and ***β***_*vm*_ by least squares and remove the fitted fixed effects to obtain residuals

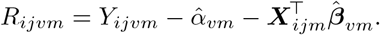

To account for batch-specific location shifts, let 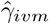 denote the fitted additive batch effect for batch *i*, feature *v*, and metric *m*. Since imaging metrics may differ substantially in scale and residual variability across features and studies, we recenter the data with respect to a batch-adjusted mean and define the standardized residuals as

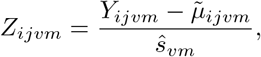

where

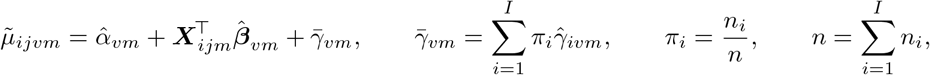

and

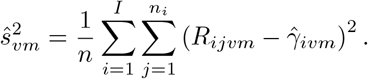

Here, *I* denotes the total number of batch levels, and 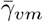 denotes either the weighted average of the fitted batch effects or the fitted effect of a prespecified reference batch. The term 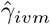 is used in 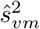 to ensure that the pooled residual variance reflects within-batch residual variation, whereas 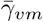 serves only as the batch-adjusted centering target for standardization. Standardization is performed separately within each feature-metric combination rather than through a single global scaling factor across all metrics. This places all metrics on a comparable working scale while preserving metric-specific scale differences for subsequent batch-effect estimation.

The standardized residuals are then stacked across metrics for each feature and subject to form the multivariate vector

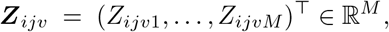

which we assume follows a multivariate normal model,

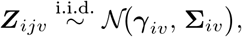

where ***γ***_*iv*_ ∈ ℝ^*M*^ is the batch-specific location effect and **Σ**_*iv*_ ∈ ℝ^*M* ×*M*^ is a symmetric positive-definite covariance matrix capturing the batch-specific dispersion across the *M* metrics for feature *v* in batch *i*, shared across subjects *j*.

The key departure from SM-ComBat is that ***Z***_*ijv*_ is modeled as a multivariate vector with a batch-specific mean and covariance, rather than as *M* independent scalars. This joint structure is what enables MM-ComBat to harmonize cross-metric covariance, not only marginal distributions.

#### 2.3.2 Estimating Batch Effect Parameters

After removing fixed effects and standardizing within each feature and metric, we model the standardized residual vectors ***Z***_*ijv*_ using two complementary Bayesian frameworks that borrow strength across features and metrics to estimate the batch-specific parameters (***γ***_*iv*_, **Σ**_*iv*_).

##### Empirical Bayes batch-effect estimation

Similar to the SM-ComBat model, we adopt an empirical Bayes framework in which hyperparameters are estimated by MoM. To ensure conjugacy, we specify multivariate normal priors on the batch-specific means and inverse–Wishart (IW) priors on the batch-specific covariances:

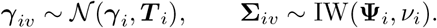

For the multivariate normal likelihood, these priors yield closed-form conditional posteriors. Using MoM plug-in estimates 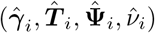, the empirical-Bayes updates are:

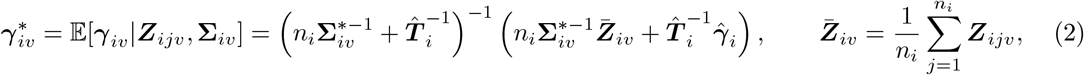

and

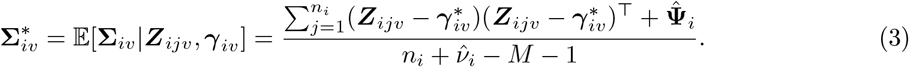

Detailed derivations are provided in Appendix A. We refer to this approach as MM-ComBat (EB). It provides a computationally efficient, closed-form solution that jointly shrinks both mean and covariance batch effects, borrowing strength across features and metrics while preserving fixed effects in the mean structure.

##### MCMC batch-effect estimation

The empirical Bayes (MoM) approach is computationally efficient and performs well for point estimation in relatively low-dimensional settings, but it has several limitations: (i) it treats hyperparameters as fixed plug-in estimates and thus underestimates uncertainty; (ii) in high-dimensional settings, the IW prior can yield unstable or biased covariance estimates, as moment matching relies only on the diagonal second moments; and (iii) the IW prior links variances and correlations through a single degrees-of-freedom parameter, inducing undesirable dependence between marginal variances and correlations.

To address these limitations, we propose a fully Bayesian framework in which fixed effects are first removed by regression and batch effects are estimated via Hamiltonian Monte Carlo (HMC) with the No-U-Turn Sampler (NUTS), a *Markov Chain Monte Carlo (MCMC)* algorithm, implemented in Stan (Hoffman and Gelman 2014; Betancourt 2017; Stan Development Team 2024; Gabry et al. 2024) (Supplementary Figure S1). Since fully joint Bayesian estimation of regression and batch-effect parameters in high-dimensional multi-metric settings would substantially increase posterior dimensionality and computational cost, leading to poor MCMC mixing and limited scalability, we adopt this two-stage strategy to allow the Bayesian model to focus on batch-related mean and covariance estimation. To increase flexibility, we replace the IW prior with an LKJ prior (Lewandowski, Kurowicka, and Joe 2009) on the correlation matrix and independent half-*t* priors for the marginal standard deviations, thereby decoupling shrinkage of variances and correlations (Barnard, McCulloch, and Meng 2000). Posterior sampling proceeds by iteratively drawing from the joint posterior distribution of (***γ***_*iv*_, **Σ**_*iv*_). After convergence, posterior means are used as point estimates for 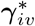 and 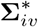. We denote this approach as MM-ComBat (MCMC). Additional details of the multivariate Bayesian model are provided in Supplementary Section S1.

#### 2.3.3 Adjusting the Data for Batch Effects

After estimating the batch-specific mean effect 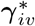 and covariance matrix 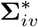, the two MM-ComBat formulations differ only in how they transform the batch-specific covariance structure.

For **MM-ComBat (baseline covariance)**, the adjusted residual vector is

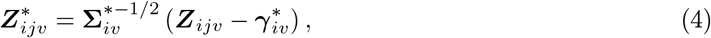

where 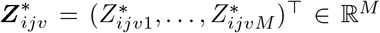. This transformation whitens the residual vector with respect to the estimated batch covariance, removing batch-induced cross-metric dependence and placing residuals on a common scale across batches. However, if the estimated batch covariance also contains biologically meaningful structure, this approach may be overly aggressive and attenuate shared biological patterns in the residuals. This is the whitening concern introduced above.

To preserve shared covariance structure, **MM-ComBat (target covariance)** maps batch-specific covariance toward a common covariance target estimated from the data:

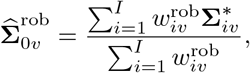

where 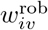 is a robust batch-specific weight. The weights are constructed in two steps. First, we form an initial weighted mean covariance

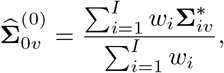

with *w*_*i*_ = *n*_*i*_*/n*, where *n* denotes the total number of observations. The Frobenius distance of each batch covariance from this initial center is then computed:

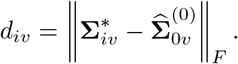

Second, we let

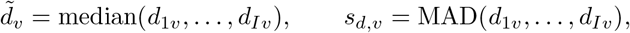

where MAD(·) denotes the median absolute deviation. We then define

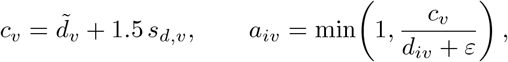

where *ε >* 0 is a small constant for numerical stability, and set

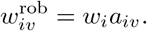

The estimator 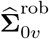 can be viewed as a robust location estimator of the shared covariance target in matrix space. This formulation is motivated by the corresponding weighted-mean estimator, which is unbiased under the idealized setting described in the Appendix C. The robust version replaces this unbiased estimator with an adaptively weighted alternative that sacrifices exact unbiasedness in exchange for greater stability and resistance to aberrant covariance estimates. Under this formulation, batch covariance estimates that remain close to the central consensus retain most of their weight, whereas unusually distant covariance estimates are downweighted. As a result, 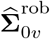 is less sensitive to outlying batch-specific covariance patterns and can provide a more stable estimate of the common covariance structure. When the number of batch levels *I* is too small (i.e., *I* ≤ 3) for reliable robust weighting, we revert to the default weighted-mean estimator to avoid instability from estimating feature-specific robustness weights from limited between-batch information.

Given the estimated target covariance, the adjusted residual vector is

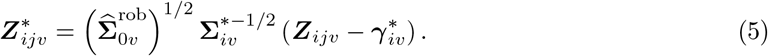

Thus, instead of whitening all residual covariance toward a standardized reference structure, this formulation maps each batch-specific covariance toward a shared target, with the goal of preserving common biological covariance patterns while removing batch-related covariance deviations.

Finally, for each metric *m*, the batch-adjusted residual component on the standardized scale is transformed back to the original scale via

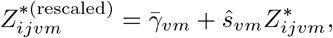

and the preserved fixed effects are added back to obtain the harmonized observation:

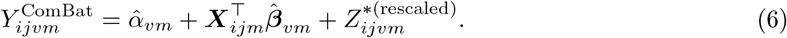

Beyond the linear-model setting, MM-ComBat can be extended within the same two-stage framework to generalized additive models (GAMs) and linear mixed-effects models, yielding multi-metric analogues of ComBat-GAM and LongComBat. In these settings, harmonization is applied to the conditional residuals obtained after fitting the corresponding mean model. For longitudinal data, the current implementation is a two-stage approximation to multivariate LongComBat. Specifically, a mixed-effects model captures within-subject dependence in the first stage, and MM-ComBat is applied to the resulting conditional residuals in the second. This is a practical working solution under the assumption that residual within-subject dependence after the first stage is weak. A fully developed multivariate extension of LongComBat with explicit longitudinal covariance modeling is left for future research.

### 2.4 Multi-metric CovBat

As discussed above, MM-ComBat assumes independence of batch effects across features and may therefore struggle to correct strong batch effects in cross-feature covariance, particularly when these interact with underlying biological variation. To address this limitation, we extend the core concept of CovBat to a multi-metric setting while preserving its two-stage harmonization framework. We refer to this approach as MM-CovBat. Intuitively, whereas MM-ComBat adjusts batch effects within each feature’s cross-metric covariance, MM-CovBat additionally targets batch effects that persist in the cross-feature covariance structure by harmonizing a shared low-dimensional latent representation learned from all metrics.

#### Stage 1 (Standard MM-ComBat for feature-wise harmonization)

We first apply MM-ComBat to remove feature-wise batch effects in the mean and covariance across metrics. This first-stage harmonization can be implemented using either the baseline covariance or target covariance formulation, depending on whether more aggressive covariance removal or better preservation of shared covariance structure is desired. For metric *m*, let **Y**^(*m*)^ ∈ ℝ^*n*×*p*^ denote the observed data matrix, and let 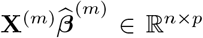 denote the fitted fixed-effects mean structure to be preserved. Since the first-stage harmonized data satisfy

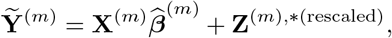

the residual matrix passed to the second stage is

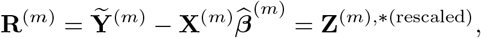

where **R**^(*m*)^ ∈ ℝ^*n*×*p*^, *n* is the number of subjects, and *p* is the number of features (e.g., ROIs).

To facilitate adjustment of batch effects in the covariance structure across features, we perform PCA on these residuals to transform them into orthogonal components, denoted as **F**^(*m*)^:

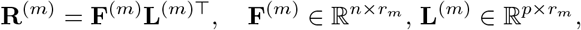

where *r*_*m*_ is the number of principal components (PCs) retained to explain most of the variation (with a default of 95%). Here **F**^(*m*)^ contains subject (score) vectors and **L**^(*m*)^ contains the corresponding feature loading vectors.

We assume that there exists a common low-dimensional latent space **G** that explains variation shared across all metric-specific PCA spaces:

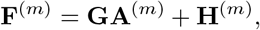

where 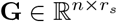 represents the shared subject scores across all metrics, *r*_*s*_ denotes the dimensionality of the shared latent subspace capturing common variation, 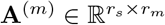 is the projection from the shared to the metric-specific subspace, and **H**^(*m*)^ is an idiosyncratic component unique to metric *m*.

#### Stage 2 (Latent space harmonization)

The main goal of the second stage is to estimate the shared latent space **G** and harmonize it to remove potential batch effects in covariance across features and, indirectly through sharing, across metrics. Without loss of generality, we impose an orthonormality constraint on **G**.

We estimate **G** and the **A**^(*m*)^ by solving

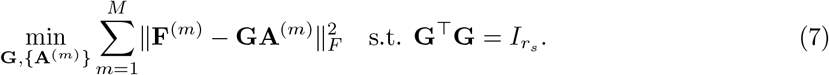

Here *M* denotes the total number of metrics, 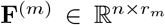 contains metric-specific PCA scores, 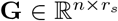 contains shared subject scores, and 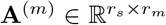 maps the shared space to the metric-specific PCA score.

Fixing **G**, the least-squares solution is **A**^(*m*)^ = **G**^⊤^**F**^(*m*)^. Let **P** = **GG**^⊤^ denote the orthogonal projection matrix onto the column space of **G**. Substituting **A**^(*m*)^ back yields:

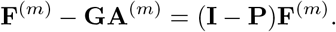

By the Pythagorean theorem, the objective function becomes:

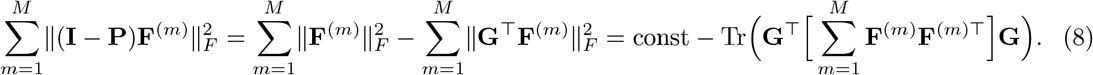

Thus, minimizing the objective is equivalent to maximizing Tr(**G**^⊤^**SG**), where 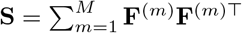. By the Rayleigh–Ritz theorem, the maximizer **G** consists of the top *r*_*s*_ eigenvectors of **S**. Detailed derivations are given in Appendix B.

We then apply standard SM-ComBat using a linear model to the columns of **G**, treating subjects as observations and latent dimensions as features, to remove batch effects in the mean and scale of the shared latent scores. The harmonized latent scores are subsequently propagated back to the metric-specific spaces via the reconstruction described above, thereby correcting covariance-related batch effects across features and metrics.

### 2.5 Simulation Design

We designed the simulation study to evaluate the proposed methods under two broad residual-covariance settings introduced above: (i) residuals contain primarily batch-related covariance and random noise, with little or no biologically meaningful covariance; and (ii) residuals contain a mixture of batch-related covariance and biologically meaningful covariance across metrics. The first setting was used to assess the batch-removal performance of the baseline-covariance formulations of MM-ComBat and MM-CovBat. The second setting was used to investigate the whitening concern for both MM-ComBat and MM-CovBat, and to compare the baseline-covariance and target-covariance formulations with respect to both batch removal and preservation of biologically meaningful covariance structure.

#### Feature-wise batch effects

We first compare MM-ComBat (baseline covariance) with SM-ComBat under two feature-wise batch-effect settings. In the *model-concordant* setting, batch-specific covariance matrices are generated from a distribution consistent with the empirical Bayes assumptions of MM-ComBat, yielding an unstructured covariance pattern aligned with the proposed model. In the *model-misspecified* setting, batch-specific covariance matrices are instead generated from a mixture of structured covariance families, including inverse-Wishart, LKJ, factor-analytic, AR, and compound-symmetry forms, thereby inducing mismatch with the exchangeable covariance prior assumed by MM-ComBat. These two settings are intended to assess the robustness of MM-ComBat under both correctly specified and misspecified covariance priors.

Within each setting, we consider three experimental conditions: two regular conditions with mild batch effects and different sample sizes, and one stress-test condition with severe batch effects. The regular conditions mimic realistic data settings, whereas the stress-test condition is designed to probe performance under extreme batch distortion.

#### Data-generating scheme

We generate biologically meaningful baseline values using three covariates: age, sex, and diagnosis. Specifically, we consider *I* = 3 batches, *p* = 70 features, *M* = 6 metrics per feature, and sample sizes *n* ∈ {100, 150, 500}. For each subject *j*, we simulate

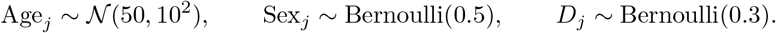

We then select a biomarker subset ℬ ⊂ {1, …, *p}* of size 0.6*p*, and allow diagnosis effects only on features in ℬ, with weaker effects in the first two metrics and stronger effects in the remaining four. The baseline values for metric *m* are generated as

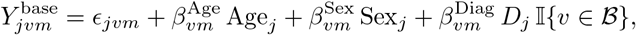

where *ϵ*_*jvm*_ represents baseline noise, and 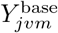 serves as the gold standard for evaluating harmonization performance, particularly the recovery of covariance structure. Under this formulation, biological signal is encoded primarily in the mean structure rather than in the cross-metric covariance.

We next introduce feature-wise batch effects. For each batch *i* and feature *v*, we first draw a location shift

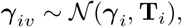

and then generate the batch-feature covariance matrix **Σ**_*iv*_ using one of two schemes: (i) an inverse-Wishart distribution in the model-concordant setting, or (ii) in the model-misspecified setting, first sampling a covariance-family label from {IW, LKJ, FA, AR, CS} with probabilities *w* = (0.20, 0.30, 0.20, 0.20, 0.10), and then drawing **Σ**_*iv*_ from the corresponding family.

Finally, for each subject *j* in batch *i*, we generate the unharmonized data as

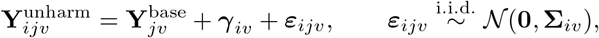

where 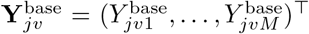 denotes the baseline cross-metric vector for subject *j* and feature *v*. This yields data with feature-wise batch effects in both the mean and covariance. The magnitudes of the covariate and batch effects vary across the three experimental conditions. Detailed parameter settings are provided in Supplementary Tables S1 and S2. For each scenario, we generate *R* = 100 independent replicates.

#### Latent-space batch effects

To further evaluate MM-CovBat and assess the robustness of MM-ComBat under structured low-rank batch variation with the baseline covariance formulation, we introduced batch effects through perturbations in a shared latent space, followed by a batch-specific transformation in the observed feature space. Specifically, we first generated a baseline latent score **G**_0*j*_ ∈ ℝ^1×*r*^ for subject *j*, where *r* denotes the dimension of the shared latent space, representing the shared latent structure underlying the observed measurements. For subjects in batch *i*, we then defined the batch-perturbed latent representation

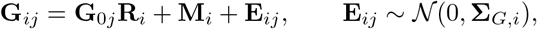

where **R**_*i*_ ∈ ℝ^*r*×*r*^ is a batch-specific latent transformation, **M**_*i*_ ∈ ℝ^1×*r*^ is a batch-specific mean-shift vector, and **E**_*ij*_ is a subject-specific latent perturbation term with batch-specific covariance **Σ**_*G,i*_, allowing the latent covariance structure to vary across batches.

For each metric *m*, the perturbed latent score **G**_*ij*_ was then mapped to the observed feature space through a metric-specific loading matrix **A**^(*m*)^, as described in the Multi-metric CovBat section. To allow these shared latent perturbations to manifest in a more realistic feature-space covariance structure, we further applied a batch-specific rotation and rescaling,

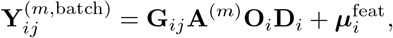

where **O**_*i*_ is a batch-specific orthogonal rotation, **D**_*i*_ is a batch-specific diagonal scaling matrix, and 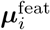 is a batch-specific feature-level mean shift. Here, 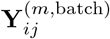 denotes the batch-induced component of the observed data for subject *j* in batch *i* and metric *m*. This additional transformation reflects the fact that low-rank technical variation may be expressed differently across observed features after projection from the shared latent space. As a result, the dominant batch structure remains shared and low-rank, while the resulting covariance distortion is allowed to vary across features in a more realistic way. This construction therefore induces structured batch effects in both the latent representation and the observed covariance structure, rather than reducing to simple feature-wise shifts or rescalings. Consequently, this setting provides a useful framework for evaluating MM-CovBat while also assessing the robustness of MM-ComBat under more complex covariance distortions.

We applied this latent batch-effect mechanism in two settings: (i) a controlled simulation based on the same data-generating framework described above, in which each scenario was repeated 200 times while varying sample size, number of features, diagnosis-effect magnitude, and the presence of additional AR(1)-correlated diagnosis effects (Controlled Simulation); and (ii) an empirical simulation based on the A2CPS dataset, restricted to a single site (UI-GE), so that controlled batch perturbations could be introduced as a data-realistic example grounded in the covariance structure of the A2CPS data (Empirical Simulation). In both settings, the simulations were designed to induce strong batch-related patterns in the residual covariance while leaving biological signals primarily in the mean structure. Together, these experiments allowed us to evaluate how effectively the competing methods removed covariance-level batch effects across features while preserving genuine biological variation in the mean structure, particularly when the standard assumption of feature-independent batch effects was violated.

#### Biological covariance patterns across metrics

Finally, to investigate the whitening concern associated with MM-ComBat (baseline covariance), we conducted an additional simulation in which the residual covariance itself contained biologically meaningful cross-metric structure. In this setting, for each feature *v*, the residual covariance was generated as

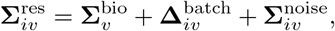

where 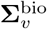 represents the biological covariance pattern shared across metrics, 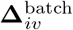 denotes a batch-specific covariance deviation, and 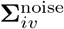 is an idiosyncratic noise component. In the current study, the biological covariance followed an AR(1) structure, while batch-related covariance deviations were generated through batch-specific orthogonal rotations of this baseline covariance. This construction allowed batch effects to alter the orientation of the covariance structure without reducing the perturbation to simple feature-wise variance shifts, thereby providing a useful setting for evaluating whether identity whitening may remove biologically meaningful covariance together with batch-related covariance variation.

Under this framework, we consider four scenarios that vary in the relative strength of biological and batch-related covariance: (1) no batch covariance with strong biological covariance, (2) moderate batch covariance with strong biological covariance, (3) moderate batch covariance with strong biological covariance and severe outliers, and (4) strong batch covariance with weak biological covariance. These scenarios are intended to distinguish settings in which identity whitening is appropriate from those in which a target-covariance formulation may better preserve biologically meaningful covariance patterns. Accordingly, in this simulation, we primarily compare MM-ComBat (baseline covariance), MM-ComBat (target covariance), and SM-ComBat. Each scenario was repeated 100 times. Details of the simulation settings are provided in Supplementary Table S4.

To further examine how the MM-CovBat framework is affected by the whitening concern, we conducted the same simulation while comparing MM-CovBat (baseline covariance), MM-CovBat (target covariance), and SM-CovBat. Here, MM-CovBat (baseline covariance) and MM-CovBat (target covariance) refer to versions of MM-CovBat that use the baseline-covariance and target-covariance formulations, respectively, in the first stage of the procedure.

Taken together, these simulations allow us to evaluate the proposed methods under both covariance-removal-oriented and covariance-preservation-oriented settings, and to characterize the trade-off between aggressive batch removal and preservation of shared biological covariance structure.

## 3 Harmonization Evaluation Methods

We evaluated harmonization performance across three complementary aspects: (1) batch removal, (2) preservation of biological signal in the mean structure, and (3) recovery of feature-wise correlation structure. For clarity, the single-metric methods, SM-ComBat and SM-CovBat, were applied separately within each metric across features, so that for each metric, the feature vector for a subject was harmonized independently of the other metrics. In contrast, the proposed multi-metric methods, MM-ComBat and MM-CovBat, were applied jointly across metrics while modeling dependence across metrics for each feature. Accordingly, comparisons between single-metric and multi-metric methods reflect the difference between harmonizing each metric separately and harmonizing all metrics jointly.

Feature-wise batch removal was assessed using univariate tests for mean differences, including ANOVA and the Kruskal–Wallis test (Kruskal and Wallis 1952), and tests for variance and scale differences, including Levene’s test (Levene et al. 1960), Bartlett’s test (Bartlett 1937), and the Fligner–Killeen test (Fligner and Killeen 1976). These univariate assessments were performed separately within each metric, yielding metric-specific results. We additionally performed multivariate assessments using MANOVA to evaluate differences in multivariate location and Box’s *M* test (BOX 1953) to assess covariance differences across batches and metrics. For these multivariate assessments, each feature was treated as a multivariate vector whose entries corresponded to the values of all metrics for that feature. All *p*-values were Bonferroni-corrected. Parametric and nonparametric alternatives are included to ensure robustness to distributional assumptions. Conclusions are drawn when results are concordant across tests within each class. Residual global batch effects were further evaluated using stratified 10-fold cross-validated random forest classifiers to predict site/scanner (R packages `caret` and `randomForest`; `ntree`=100, all other hyperparameters set to their default values), with the macro-averaged area under the ROC curve (AUC) used to quantify residual batch-related variation.

To evaluate preservation of biologically meaningful signal in the mean structure, we fit feature-wise regression models using prespecified covariates and counted the number of significant features. For the A2CPS dataset, where the true biological effects are unknown, we controlled the false discovery rate (FDR) using the Benjamini–Hochberg (BH) procedure. In simulation studies, where the truth was known, we reported the number of true-positive discoveries and the empirical FDR to assess how accurately each method recovered true associations.

We evaluated correlation recovery by comparing within-metric and cross-metric (pooled-feature) correlation structures with the ground truth in simulation studies. Similarity between the estimated and true correlation matrices was quantified using four matrix-distance metrics: the Frobenius norm and element-wise mean squared error (MSE) to assess overall recovery, and eigenvalue error and spectral norm to detect structural distortion. Frobenius norm and MSE capture average element-wise recovery, while spectral norm reflects worst-case directional distortion and eigenvalue error quantifies structural deformation of the correlation spectrum. Lower values indicate better recovery. These same metrics were also used to assess batch-related differences in correlation matrices across batch levels, with smaller values indicating weaker residual batch effects.

For the EB approach, we additionally evaluated model assumptions by comparing the empirical distributions of estimated additive batch effects with their corresponding prior distributions. For multiplicative effects, we performed prior predictive checks by comparing empirical covariance matrices with draws simulated from the inverse-Wishart prior. Matrix-distance metrics were used to quantify deviations from the prior mean, and histograms of these statistics were used to visualize agreement between the empirical and prior distributions. Greater overlap indicated that the EB assumptions were reasonably satisfied. For the fully Bayesian approach, convergence of the Markov chains was assessed using standard diagnostics, including the potential scale reduction factor 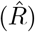, effective sample size, and visual inspection of trace plots, with 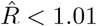 used as the convergence criterion.

To more directly assess the trade-off between batch removal and preservation of biologically meaningful residual covariance under the whitening concern (evaluated in the biological covariance simulation of Section 2.5), we introduce three additional criteria specific to this setting. Each targets a distinct aspect of the baseline-versus-target-covariance trade-off. (i) *Biological covariance error:* the relative Frobenius error between the estimated and true biological covariance matrices, quantifying how much meaningful covariance structure is distorted by harmonization; (ii) *batch covariance heterogeneity:* the average relative pairwise Frobenius distance between batch-specific covariance matrices, measuring how much residual batch-related covariance variation remains after harmonization; and (iii) *batch AUC:* the site-prediction from cross-validated random-forest classification of batch labels, consistent with the global batch-detection metric used throughout. For all three criteria, lower values are preferred as lower biological covariance error and batch covariance heterogeneity indicate better covariance preservation and batch removal respectively, while a batch AUC near 0.5 indicates batches are no longer distinguishable. Together, these criteria allow direct comparison of how aggressively each formulation removes covariance-level batch effects relative to how much biological covariance structure it retains.

## 4 Results

### 4.1 A2CPS Study

In the A2CPS study, we considered age, sex, and pre-surgical pain duration as biologically meaningful sources of variation to be preserved during harmonization. Although site-level differences were observed, particularly in pain duration, these covariates showed sufficient cross-site overlap for inclusion in the harmonization model, enabling adjustment for site/scanner effects while preserving relevant participant-level variation and reducing potential confounding (Table 1).

**Table 1.** Demographic characteristics of A2CPS participants stratified by site.

| Variable | Overall | WS | UI | UM | SH | UC | NS |
| --- | --- | --- | --- | --- | --- | --- | --- |
| **Number of participants** | 473 | 29 | 130 | 97 | 22 | 82 | 113 |
| **Sex** |  |  |  |  |  |  |  |
| Male | 161<br>(34.04%) | 10<br>(34.48%) | 34<br>(26.15%) | 47<br>(48.45%) | 8<br>(36.36%) | 21<br>(25.61%) | 41<br>(36.28%) |
| Female | 312<br>(65.96%) | 19<br>(65.52%) | 96<br>(73.85%) | 50<br>(51.55%) | 14<br>(63.64%) | 61<br>(74.39%) | 72<br>(63.72%) |
| **Age** | 63.38<br>(9.68) | 61.00<br>(10.07) | 62.25<br>(8.91) | 61.86<br>(12.05) | 60.45<br>(11.01) | 63.78<br>(9.43) | 66.88<br>(6.91) |
| **Pain duration** | 53.01<br>(86.88) | 43.86<br>(64.87) | 54.96<br>(89.83) | 39.40<br>(65.31) | 9.82<br>(25.45) | 63.72<br>(98.17) | 65.42<br>(99.60) |

*Note.* Values are presented as sample size, number with percentage underneath for sex, and mean with standard deviation underneath for age and pre-surgical pain duration. Percentages for sex were calculated within each site. Although site-level differences were observed, particularly in pain duration, the covariates showed sufficient cross-site overlap for inclusion in the harmonization model.

#### Regression Model Specification

Exploratory plots of cortical-region metrics versus age and pain duration suggested nonlinear associations (Figure 1A,B). We therefore fit generalized additive models (GAMs) to estimate fixed effects to be preserved during harmonization. Sex was included as a parametric term, while age and pre-surgical pain duration were modeled as smooth functions. As shown in Figure 1C, strong residual cross-metric correlations persisted among some metrics, and the correlation patterns differed across batch levels, suggesting scanner-related effects not captured by the initial model and retained in the cross-metric covariance. Residual distributions, stratified by site, indicated both additive (mean-shift) and multiplicative (scale) batch effects across all cortical metrics, with magnitudes varying by ROI and metric. We also observed influential outliers in some metrics, such as FoldInd, consistent with sensitivity of folding index to local surface irregularities and noise amplification, which may pose challenges for ComBat-type methods (Figure 1D).

**Figure 1.**
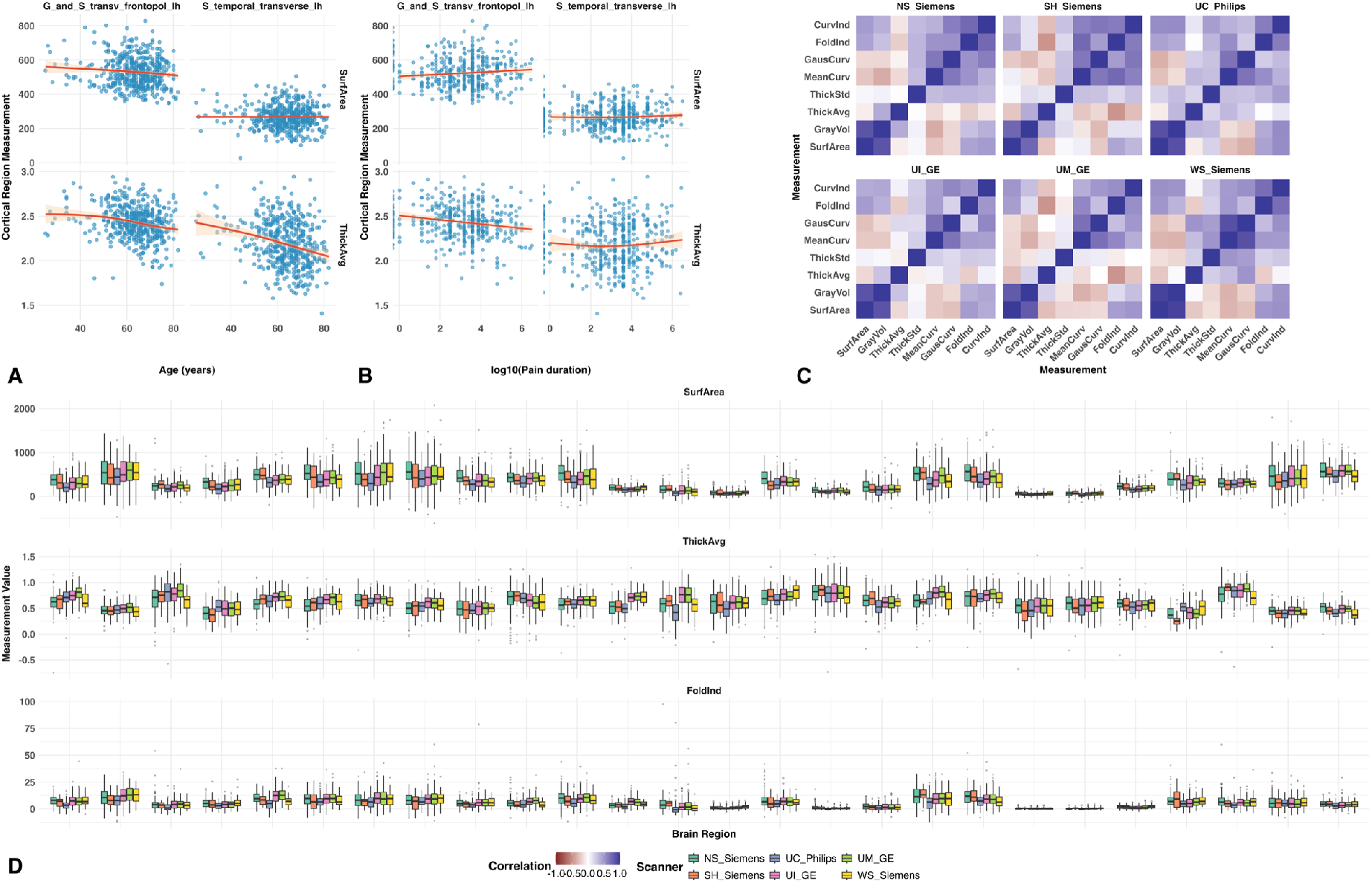
Exploratory Analysis. (A–B) Scatterplots of cortical-region metrics against age and pre-surgical pain duration show nonlinear associations, motivating the use of generalized additive models (GAMs) for covariate adjustment. (C) Residual cross-metric correlation matrices, stratified by batch, show strong correlations among some metrics and distinct correlation patterns across sites, consistent with scanner-related effects not captured by fixed effects. (D) Residual distributions by site, highlighting additive and multiplicative batch effects across regions and metrics. Some metrics (e.g., FoldInd) exhibit influential outliers in the underlying data, which affect the scale.

#### Feature-wise Batch Detection

We compared five harmonization methods: (1) MM-ComBat (EB), (2) MM-ComBat (MCMC), (3) SM-ComBat, (4) MM-CovBat, and (5) SM-CovBat, using GAMs to estimate fixed effects. In the A2CPS application, the EB implementation of the multi-metric framework required approximately 16 seconds, compared with 13 seconds for the single-metric approach, whereas the MCMC implementation required roughly 13 hours. These results indicate that the EB version has a computational cost similar to that of the single-metric approach, while the MCMC version is substantially more computationally intensive. For simplicity, only the EB variant was applied to both CovBat methods. For MM-ComBat and MM-CovBat, we focused on the baseline covariance approach, while results for the target covariance approach are provided in the Supplementary Material as a sensitivity analysis. Overall, the EB assumptions for MM-ComBat (EB) were well met for this dataset (Supplementary Figure S2). Additionally, for the MCMC model, all parameters exhibited 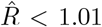 and effective sample sizes exceeding 400, indicating satisfactory convergence to the target posterior distribution. Feature-wise batch-effect diagnostic results were then compared across the five harmonization methods as well as the unharmonized data.

Univariate test results (Figure 2A) showed that all harmonization methods successfully removed additive batch effects within each metric, whereas most methods had more difficulty fully eliminating scaling batch effects in some metrics. Focusing on these metric-specific scaling effects, both MM-ComBat methods outperformed SM-ComBat for MeanCurv and GausCurv, but not for CurvInd and FoldInd, likely reflecting the influence of outliers. Between the two MM-ComBat variants, the EB approach performed better for MeanCurv, whereas the MCMC approach performed better for GausCurv and CurvInd. However, the maximum difference in the univariate test results was only about 2%, suggesting that all harmonization methods achieved generally good feature-wise batch removal in mean and variance within each metric. Multivariate tests (Figure 2B) revealed that SM-ComBat failed to adequately remove batch effects in covariance across metrics, with 70% of ROIs showing significant differences according to Box’s M test. In contrast, both MM-ComBat methods substantially reduced these effects, and the MCMC variant left no ROIs with significant covariance differences after multiple-testing correction. For differences in multivariate mean structure across metrics, MANOVA showed no remaining significant batch effects after harmonization, indicating that all methods were effective in removing cross-metric batch-related shifts in location. This is consistent with the univariate ANOVA results, which likewise suggested good control of additive batch effects within each metric. Comparing the ComBat and CovBat frameworks in both the multi-metric and single-metric settings, we found that their performances were broadly similar in both the univariate and multivariate analyses. Together, these results indicate that joint modeling substantially improves feature-wise cross-metric covariance harmonization.

**Figure 2.**
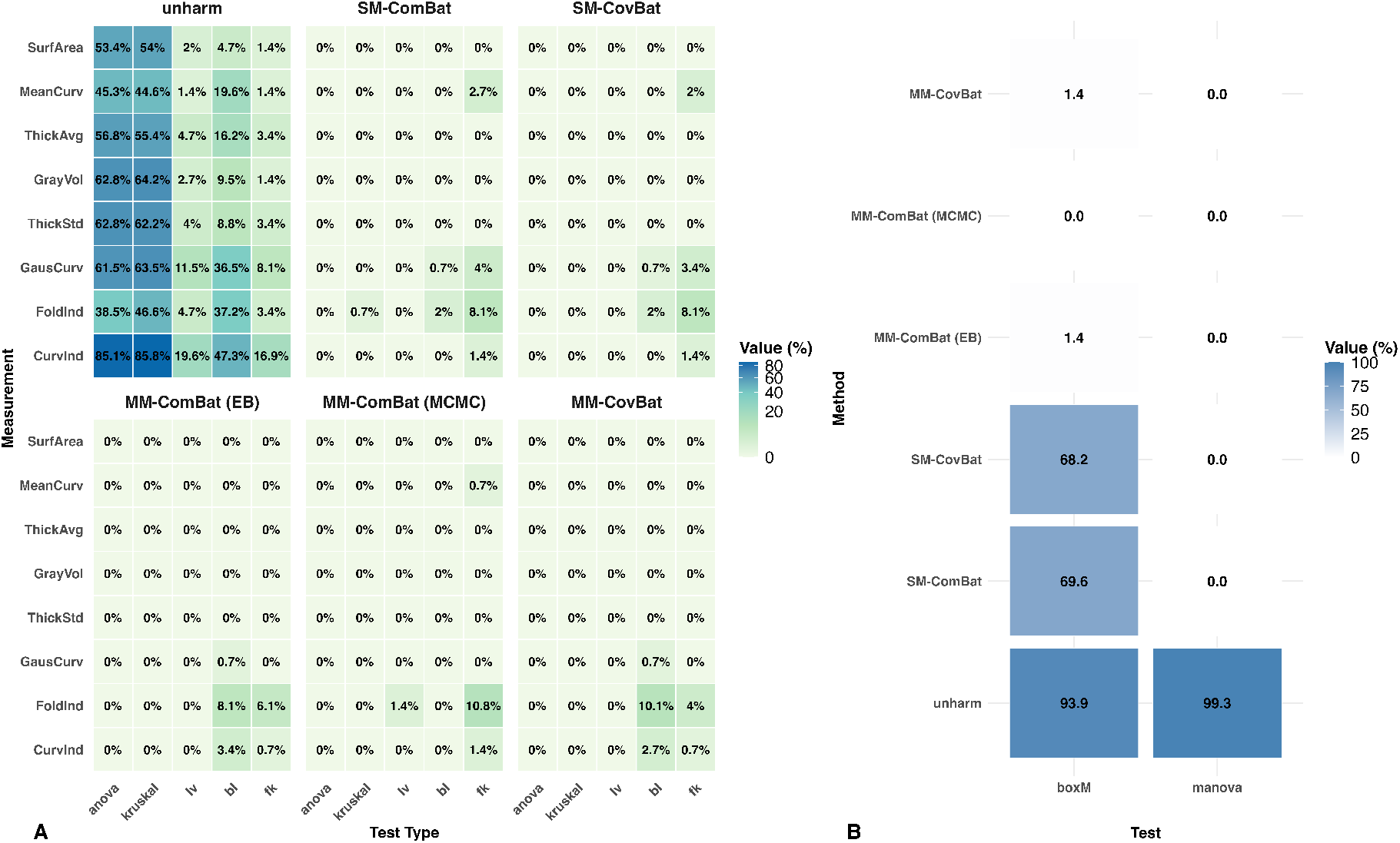
Statistical Tests. (A) Univariate statistical tests within each metric comparing (1) unharmonized data, (2) SM-ComBat, (3) SM-CovBat, (4) MM-ComBat (EB), (5) MM-ComBat (MCMC) and (6) MM-CovBat with GAM-based fixed effects preserved. Additive effects are tested using ANOVA and Kruskal–Wallis; multiplicative effects using Levene’s, Bartlett’s, and Fligner–Killeen. Both MM-ComBat variants outperform SM-ComBat for MeanCurv and GausCurv, but not for CurvInd and FoldInd where influential outliers limit gains. Within MM-ComBat, EB slightly exceeds MCMC for MeanCurv, while MCMC is better for GausCurv and CurvInd. (B) Multivariate statistical tests of location and covariance across metrics (MANOVA and Box’s M). SM-ComBat leaves substantial covariance differences, whereas MM-ComBat markedly reduces them. In particular, MM-ComBat (MCMC) effectively removes covariance batch effects. SM-CovBat slightly outperformed SM-ComBat.

#### Fixed-effects Preservation and Global Batch Detection

We next assessed whether harmonization preserved nonlinear covariate effects using GAMs and whether residual global batch effects remained within each metric. Age effects were consistently strong, while pain duration effects were considerably weaker. Notably, in GausCurv, the unharmonized dataset yielded more significant findings, some of which may reflect residual batch effects. That is, a decrease in the number of significant findings does not necessarily imply a loss of true signal. After controlling FDR, MM-ComBat generally retained as many or more signals than SM-ComBat, and substantially more signals for SurfArea (Figure 3A). We therefore conclude that MM-ComBat preserved more putative biological signals overall.

**Figure 3.**
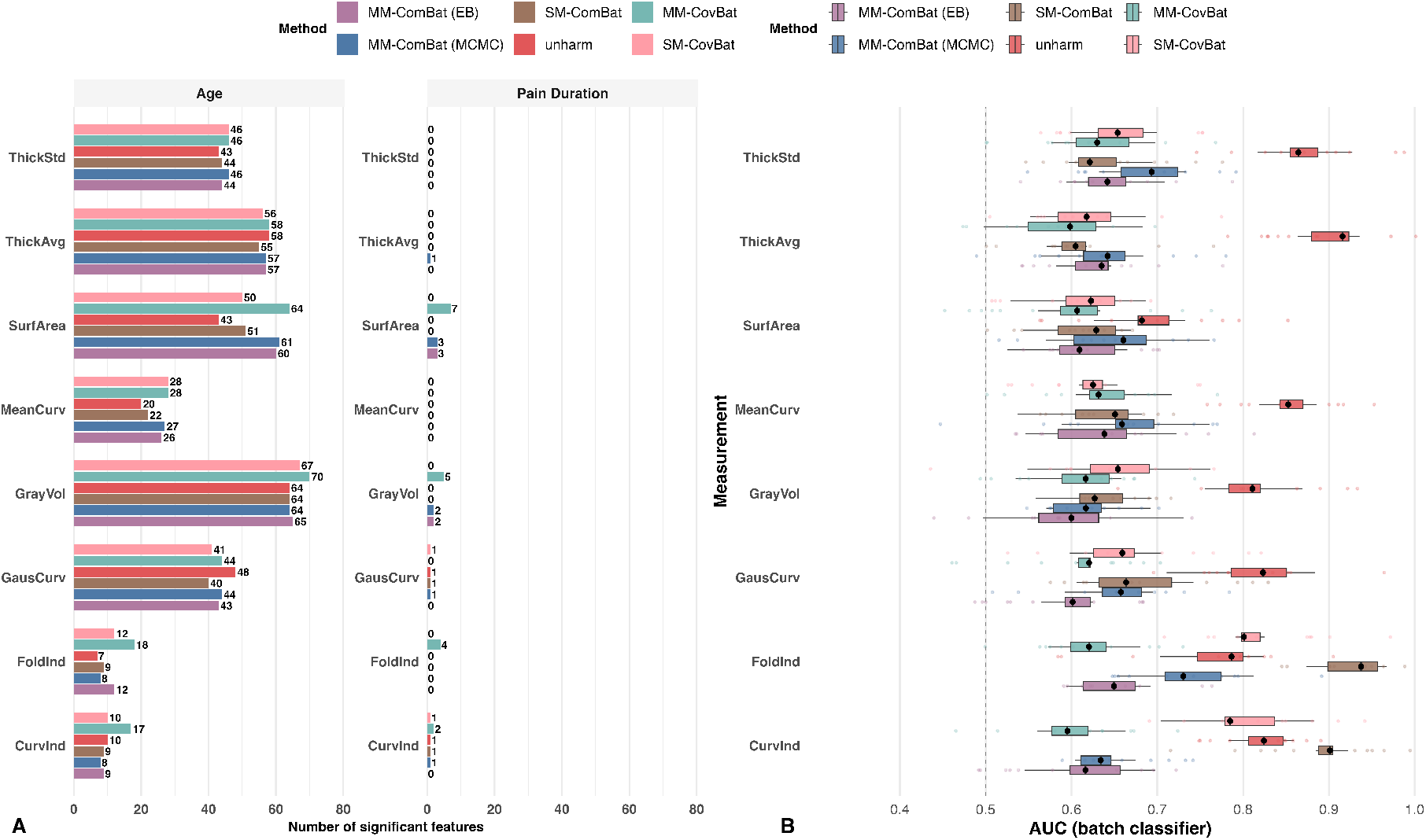
Detection of Batch Effects and Biological Variation. (A) Number of ROIs with significant nonlinear fixed effects from GAMs fit to unharmonized data and to data harmonized by MM-ComBat (EB), MM-ComBat (MCMC), SM-ComBat, MM-CovBat, and SM-CovBat. Across most metrics, MM-ComBat preserves at least as many signals as SM-ComBat, with substantial gains for SurfArea. Pain duration shows weaker effects overall. Both SM-CovBat and MM-CovBat identified more significant signals than their ComBat counterparts, with MM-CovBat detecting substantially more. (B) Residual batch signal measured by 10-fold CV random-forest AUC for predicting site/scanner from features. Lower AUC indicates better batch removal. MM-CovBat yields the lowest AUCs for most metrics. SM-CovBat outperforms SM-ComBat but still shows high AUCs for CurvInd and FoldInd, indicating poor batch removal likely driven by severe outliers. MM-ComBat (MCMC) is slightly less effective than MM-ComBat (EB) in most metrics, but generally outperforms SM-ComBat and is more robust when outliers are present.

Evaluating global batch signals within each metric, the random-forest classifier showed that SM-ComBat yielded slightly lower AUCs for ThickStd and ThickAvg, but most values were below 0.7 for all methods, indicating weak residual batch signals. For other metrics, MM-ComBat (EB) performed best among the ComBat methods, whereas SM-ComBat exceeded 0.9 for CurvInd and FoldInd, reflecting its failure to remove batch effects in the presence of severe outliers (Figure 3B). MM-ComBat (MCMC) was slightly less effective than the EB variant in most metrics, likely due to prior misspecification and cross-metric borrowing, but still outperformed SM-ComBat, particularly in outlier-prone metrics.

Extending to covariance harmonization, both SM-CovBat and MM-CovBat detected more significant signals than their ComBat counterparts, with MM-CovBat showing the greatest gains. Although SM-CovBat outperformed SM-ComBat in outlier-prone metrics, the AUCs for SM-CovBat remained high (around 0.8), indicating that considerable batch effects persisted. Compared with MM-ComBat (EB), MM-CovBat further improved correction in FoldInd, CurvInd, ThickAvg, and ThickStd, achieving the best performance in global batch removal. Taken together, these results indicate that the CovBat frameworks outperformed the ComBat frameworks in removing residual global batch signals present in feature values and their correlations at the metric level, thereby improving the detection of biologically meaningful signals, with MM-CovBat achieving the strongest overall performance.

#### Batch Detection in Correlation

Finally, we evaluated batch removal in correlations by comparing the average distance between pairwise correlation matrices across batches using the four previously described distance metrics. SM-CovBat performed best for within-metric correlations, which is consistent with its single-metric harmonization design. MM-CovBat outperformed MM-ComBat in all metrics except the spectral norm (Figure 4A), indicating strong correction with minimal structural distortion. For cross-metric correlations, MM-CovBat achieved the most effective batch removal (Figure 4B), highlighting its advantage in jointly modeling batch effects across metrics and performing latent space harmonization.

**Figure 4.**
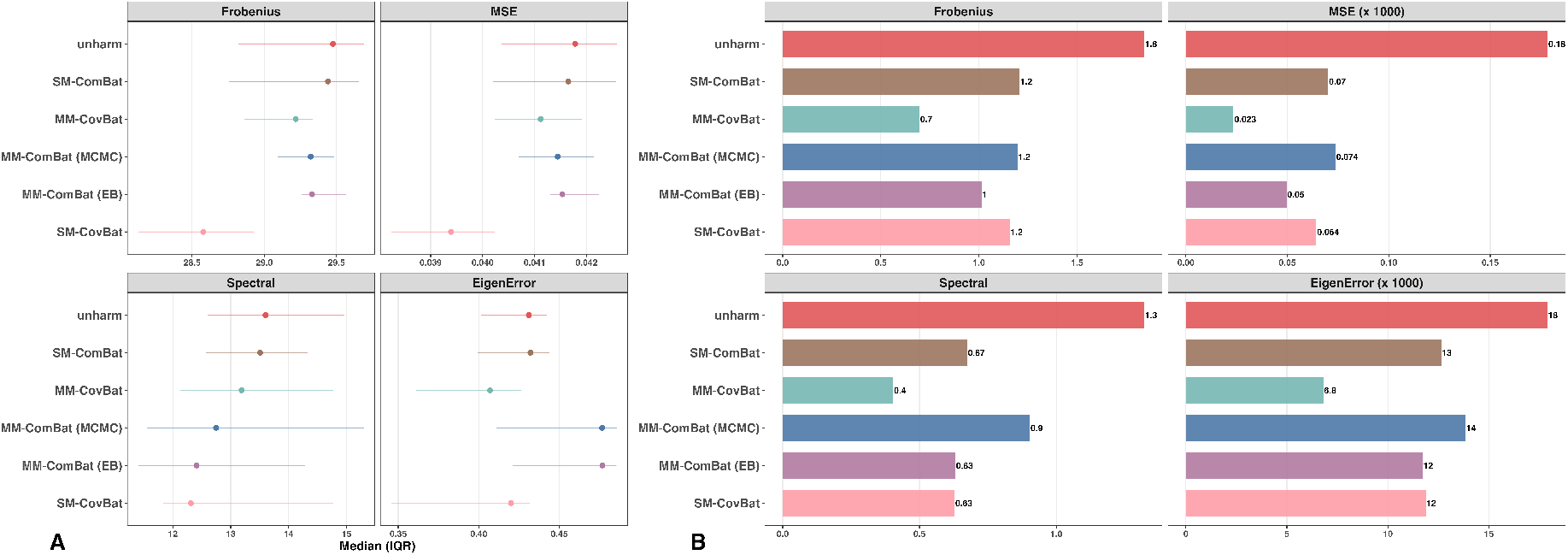
Batch Detection in Correlation. (A) Average within-metric correlation matrix distance across batch levels. Points indicate medians across ROIs, and horizontal error bars denote the interquartile range (IQR; 25th–75th percentiles). SM-CovBat performs best overall, while MM-CovBat outperforms both MM-ComBat variants. (B) Average cross-metric correlation matrix distance across batch levels. MM-CovBat performs best overall. For visualization, MSE (×1000) and EigenError (×1000) indicate that values were multiplied by 1000 to improve scale comparability.

### 4.2 Simulation

#### Feature-wise Batch-effect Simulation

To better understand the performance of MM-ComBat in removing feature-wise batch effects across metrics, we applied (i) MM-ComBat (EB), (ii) MM-ComBat (MCMC), and (iii) SM-ComBat to simulated data, preserving age, sex, and diagnosis effects using linear models. As noted above, both MM-ComBat methods were implemented using the baseline covariance formulation. Harmonization performance was evaluated under three experimental conditions within the model-concordant scenario, with additional results under model misspecification reported to assess the robustness of the proposed methods.

##### Batch removal

Following the A2CPS study, we assessed within-metric global batch effects using 10-fold cross-validated random forest AUC. All three methods substantially reduced batch effects across the three experimental conditions in both scenarios, with mean AUCs for batch detection remaining around 0.6 for smaller sample sizes and further decreasing to 0.56 for the largest sample size (n = 500) (Table 2; Supplementary Table S5). Accordingly, within-metric AUCs were similar across methods. This is expected because the evaluation focuses on within-metric marginal batch effects, a setting in which SM-ComBat is specifically designed to perform well and therefore performs comparably to MM-ComBat in this global batch-effect setting.

**Table 2.**
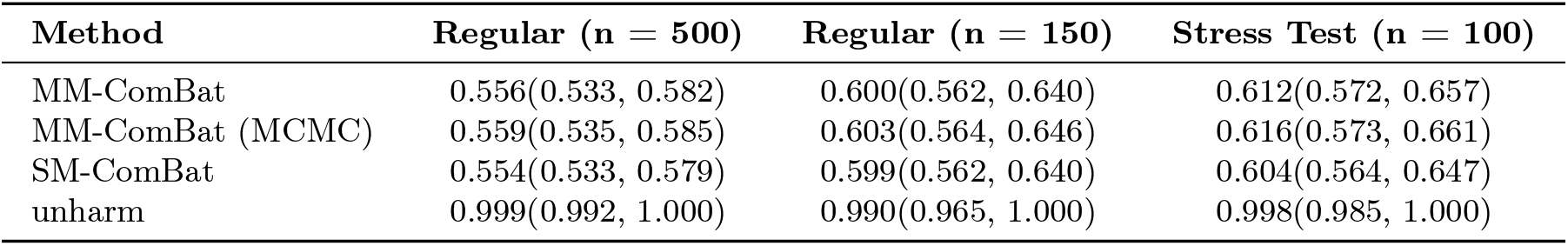
Within-metric batch-effect removal performance across experimental conditions (point estimates with 95% confidence intervals) under the model-concordant scenario.

In Box’s M test, under the model-concordant scenario, about 25% and 60% of features in the unharmonized dataset showed covariance-related batch effects in the regular conditions with sample sizes of 150 and 500, respectively, and roughly 30% in the stress test. After SM-ComBat, these proportions decreased to approximately 12% and 37% in the regular conditions but increased to 50% in the stress test. In contrast, both MM-ComBat variants produced datasets largely free of covariance-related batch effects, with the MCMC variant performing particularly well (Figure 5A). Under the model-misspecified scenario, results were similar, except that MCMC remained robust under mild batch effects but was occasionally less stable in the stress test due to prior mismatch (Figure S3A).

**Figure 5.**
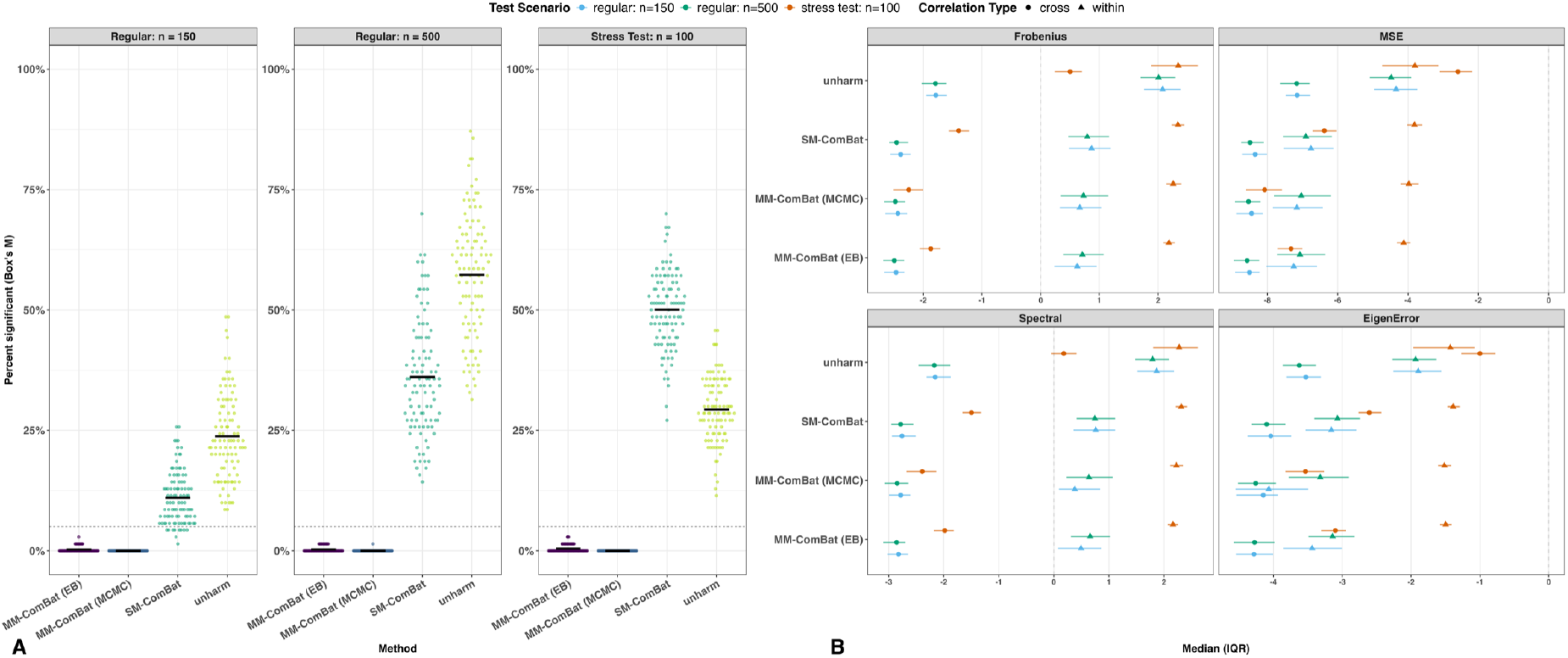
Batch Detection in Covariance and Correlation Recovery. (A) Box’s M test across experimental conditions. Both MM-ComBat variants outperformed SM-ComBat, with the MCMC variant showing greater gains in smaller samples. (B) Correlation recovery. The MCMC approach achieved lower eigenvalue error for within-metric correlations under regular conditions with small samples and better cross-metric recovery under stress when priors were correctly specified (for misspecification results, see Supplementary Figure S3).

Overall, these findings highlight a key strength of MM-ComBat: its ability to mitigate batch effects in cross-metric covariance. By comparison, SM-ComBat effectively removed within-metric batch effects but struggled to address covariance-related effects across metrics.

##### Correlation Recovery

Under the model-concordant scenario, cross-metric correlations generally showed smaller deviations from the gold standard than within-metric correlations, reflecting the clean, homoscedastic structure of the simulated data (Figure 5B). For within-metric recovery, both MM-ComBat variants outperformed SM-ComBat across all experimental conditions. The MCMC variant showed substantially better eigenvalue error performance at a sample size of 150, although this advantage diminished with larger samples. In the stress test, all methods performed similarly, with the EB variant showing slightly better performance, potentially indicating greater robustness. For cross-metric correlations, both MM-ComBat variants markedly outperformed SM-ComBat in the stress test, with the MCMC variant performing best.

Under model misspecification (Supplementary Figure S3B), results were largely consistent with the model-concordant scenario. However, the advantage of the MCMC variant over the EB variant for cross-metric recovery in the stress test disappeared, likely due to prior misspecification. Overall, MM-ComBat (MCMC) exhibited stronger performance in preventing within-metric correlation distortion (especially in small samples) and in recovering cross-metric correlations when batch effects were extremely strong and priors were correctly specified. The MCMC approach was robust to prior misspecification under mild batch effects, whereas the EB variant was more robust to model misspecification.

##### Signal Preservation

We fitted linear models to assess age, sex, and diagnosis effects. As designed, all ROIs showed strong age effects and moderate-to-strong sex effects, while 60% served as biomarkers with weak diagnosis effects in the first two metrics and moderate-to-strong effects in the remaining ones. We focused on weak and moderate-to-strong signals, as these are more likely to be removed along with batch effects under confounding. MM-ComBat (MCMC) generally preserved the most fixed effects while maintaining FDRs comparable to SM-ComBat. Its advantages were most pronounced in smaller samples, under the stress test, and for moderate-to-strong signals (Figure 6A). FDR decreased with increasing sample size and stronger signals (Figure 6B). In the stress test, SM-ComBat and unharmonized data exhibited lower mean FDRs due to missed detections that artificially reduced averages. Thus, the slightly higher FDRs observed with MM-ComBat should not raise concern, as they likely reflect increased sensitivity rather than inflated false positives. Under model misspecification, results were similar, though the MCMC advantage over EB diminished due to prior misspecification (Supplementary Figure S4). Overall, MM-ComBat achieved superior preservation of moderate-to-strong biological signals.

**Figure 6.**
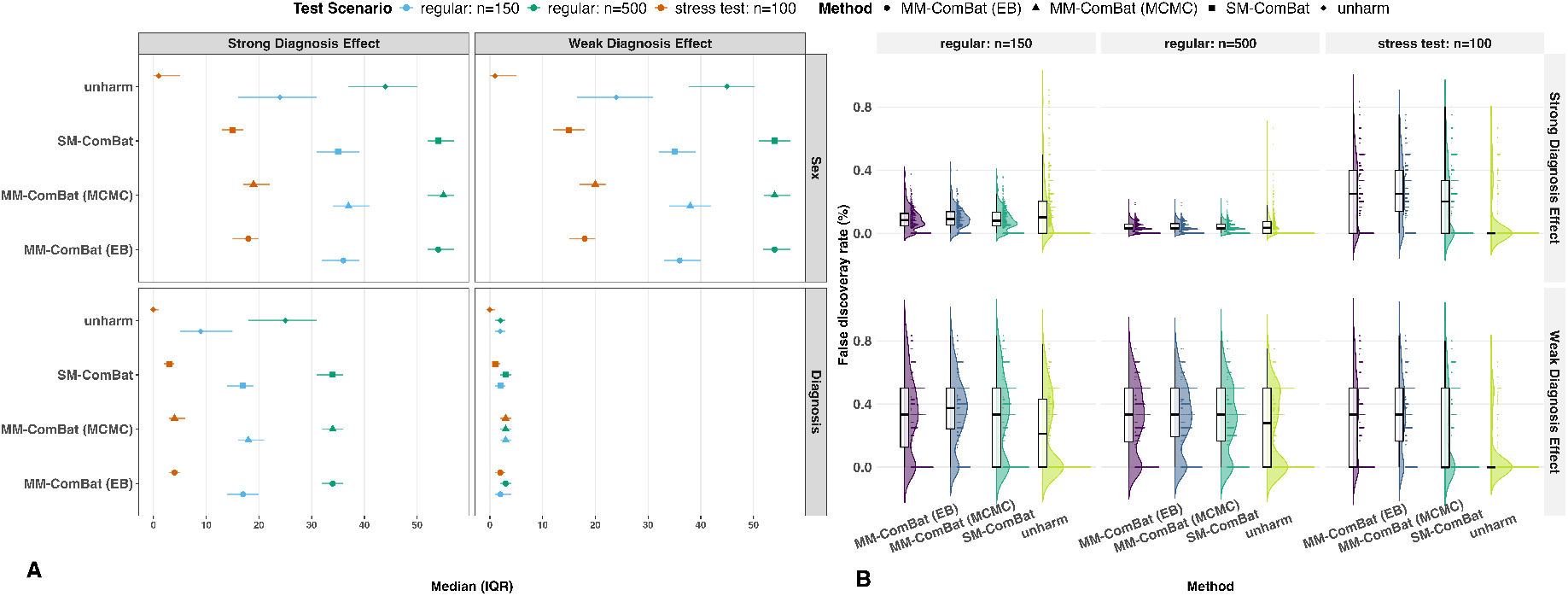
Fixed-effects Detection and Corresponding FDRs. (A) Preservation of biological signals. Counts of features with significant sex and diagnosis effects across experimental conditions. MM-ComBat (MCMC) generally preserved the most fixed effects, with greater advantages in smaller samples, under stress, and for moderate-to-strong signals. (B) False discovery rate (FDR) based on known biomarkers. Both MM-ComBat variants performed comparably to SM-ComBat, although SM-ComBat showed lower mean FDRs in the stress test due to missed detections rather than improved specificity (for misspecification results, see Supplementary Figure S4).

#### Latent-space Batch-effect Simulation

As noted earlier, MM-ComBat assumes independent batch effects across features and may therefore struggle when strong batch effects remain in feature correlations. To evaluate whether MM-CovBat addresses this limitation, we applied: (1) MM-ComBat, (2) MM-CovBat, (3) SM-ComBat, and (4) SM-CovBat (all using the EB approach) to the Controlled Simulation (linear model preserving age, sex, and diagnosis) and to the Empirical Simulation (GAM preserving age, sex, and pain duration). As before, both MM-ComBat and MM-CovBat were implemented using the baseline covariance formulation. In this section, we focus on two key aspects: (1) removal of batch effects in feature correlations and (2) preservation of fixed effects measured by FDR, as these represent the main advantages of the CovBat framework.

##### Controlled Simulation

We calculated the average pairwise distances between correlation matrices across batch levels for both within- and cross-metric correlations using the four matrix-distance criteria. For within-metric correlations, MM-CovBat consistently achieved the smallest distances across all criteria, particularly in smaller samples with more features (Figure 7A). As sample size increased, the gap between MM-CovBat and MM-ComBat narrowed. Both multivariate methods (MM-ComBat and MM-CovBat) outperformed their univariate counterparts across all sample sizes and feature counts.

**Figure 7.**
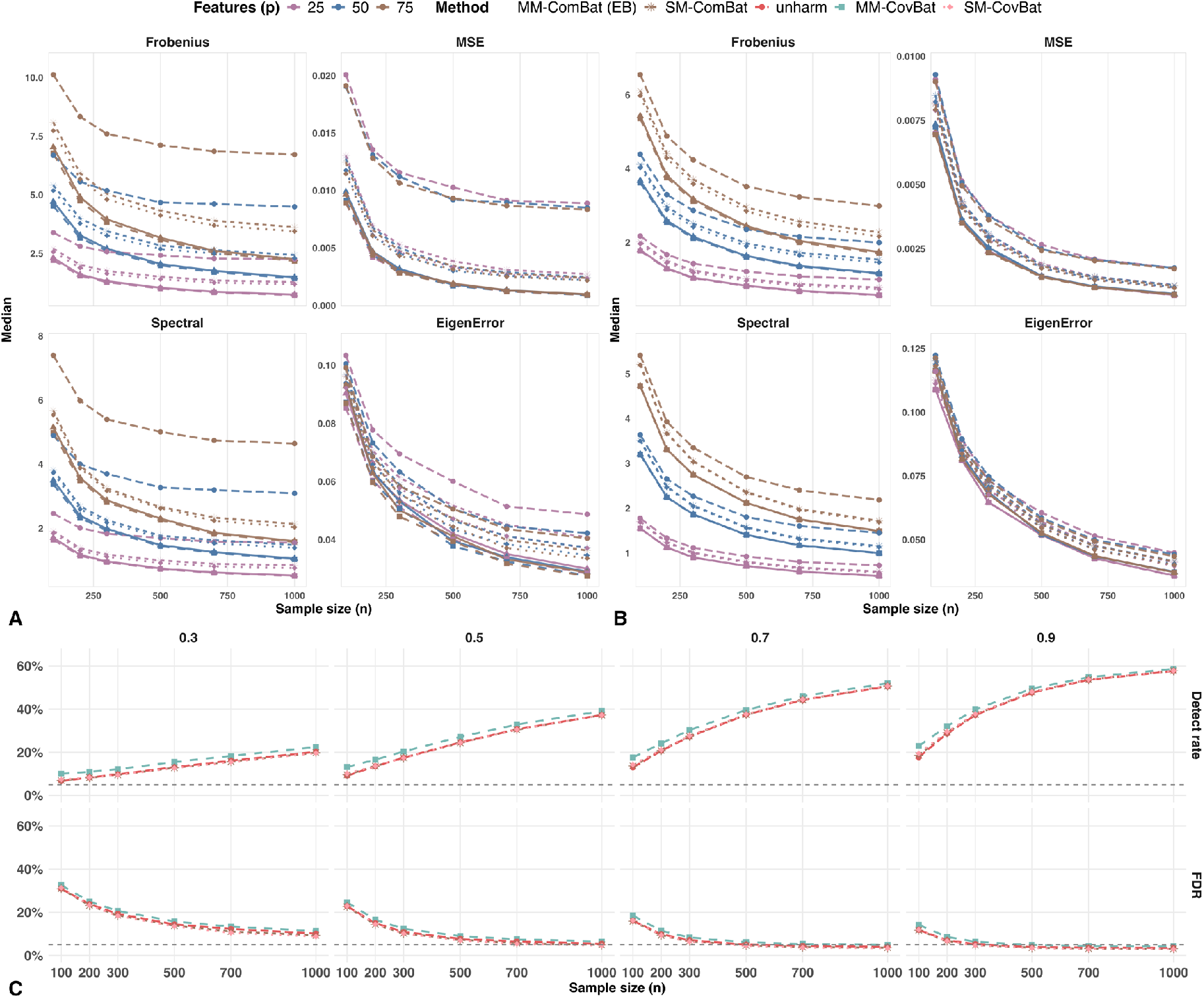
Batch Detection in Correlation and Diagnosis Effect Preservation. (A) Average within-metric correlation matrix distance across batch levels. Both MM-CovBat and MM-ComBat outperform SM-CovBat and SM-ComBat, with MM-CovBat showing greater improvement, particularly in small sample sizes and large feature sets. (B) Average cross-metric correlation matrix distance across batch levels. MM-CovBat continues to perform best, with minimal differences from MM-ComBat. (C) Detection of diagnosis effects and corresponding FDRs. MM-CovBat consistently identifies more true signals, with slightly higher FDRs. When the diagnosis effect size is 0.9 and the sample size is 500, the FDR for MM-CovBat drops below 0.05.

For cross-metric correlations, we observed a similar pattern, though the advantage of MM-CovBat over MM-ComBat was smaller (Figure 7B), consistent with the simulation design in which cross-metric batch effects were relatively weak and homogeneous across batches. In terms of fixed-effect preservation, MM-CovBat consistently detected more true signals, accompanied by slightly higher FDRs (Figure 7C), reflecting the common power–FDR trade-off. Differences in FDR across methods were minimal, and as sample size and effect size increased, all methods detected more diagnosis effects while maintaining lower FDRs. With moderate-to-strong diagnosis effects (*β*_*d*_ = 0.9) and a median sample size (*n* = 500), the FDR for MM-CovBat remained below 0.05. These results highlight MM-CovBat’s strength in preserving genuine biological signals, likely facilitated by improved correction of correlation-related batch effects.

##### Empirical Simulation

We randomly assigned three batch levels to the single-site subset of the A2CPS data (130 observations) and introduced latent batch effects following the same procedure as in the controlled simulations. MM-ComBat performed comparably to or slightly better than MM-CovBat in removing within-metric batch effects, likely due to its greater robustness to outliers (Figure 8A). Both methods, however, substantially outperformed SM-CovBat and SM-ComBat. The PCA plot of the **G** space (Figure 8C) revealed potential confounding between batch levels and shared latent scores after SM-CovBat, as indicated by pronounced directional variation within clusters. Both MM-ComBat and MM-CovBat greatly reduced these directional differences in the first two principal components, with MM-CovBat performing better in the higher-order components. Moreover, MM-CovBat achieved superior performance in pooled correlations across metrics for all four criteria (Figure 8B), highlighting its strength in correcting cross-feature and cross-metric covariance distortions that arise under strong multivariate batch effects.

**Figure 8.**
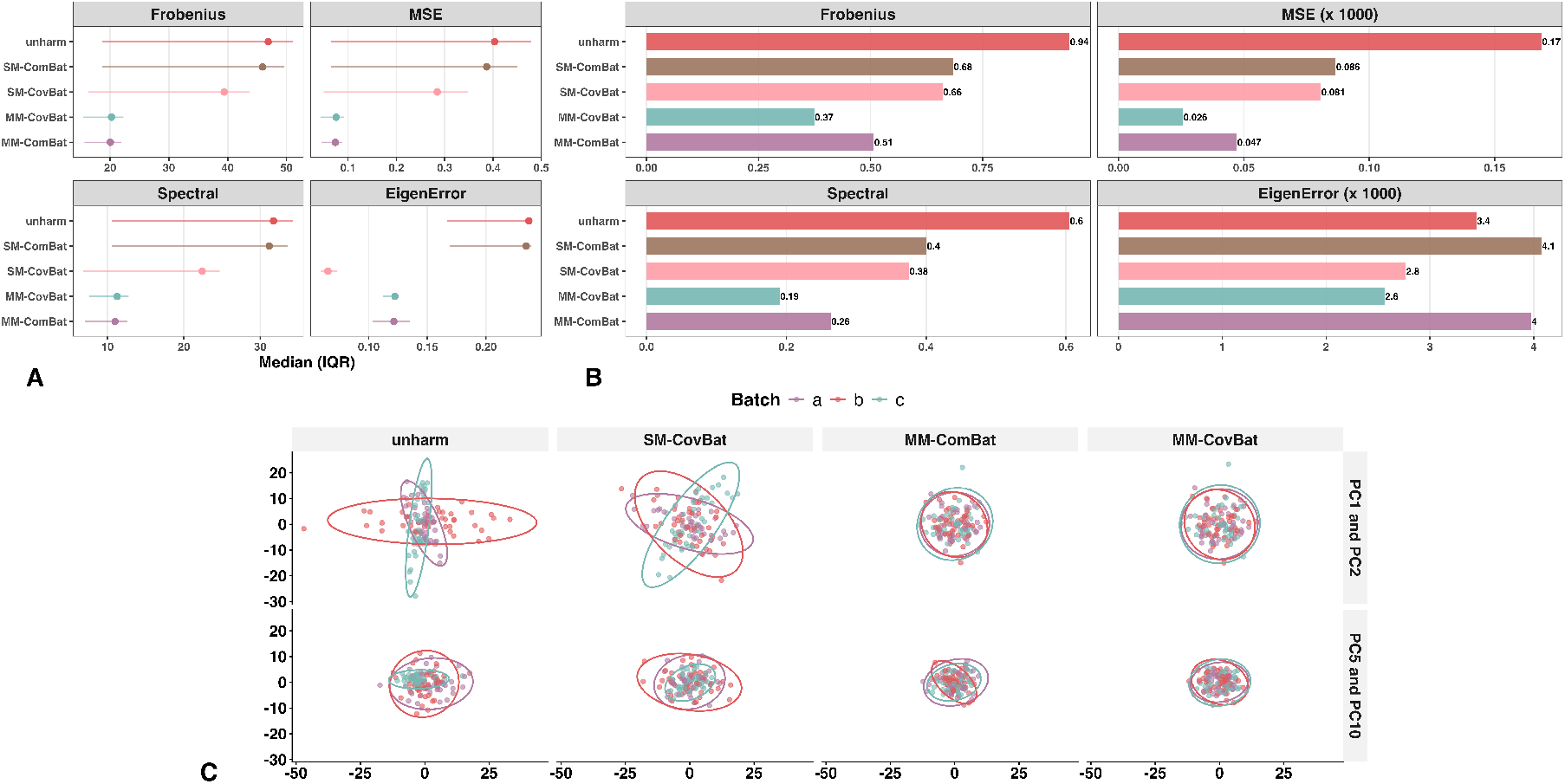
Batch Detection in Correlation. (A) Average within-metric correlation matrix distance across batch levels. MM-CovBat performs comparably to MM-ComBat and substantially better than both univariate methods. (B) Average cross-metric correlation matrix distance across batch levels. MM-CovBat shows the best overall performance. (C) PCA of the **G** space. Residuals after SM-CovBat display pronounced directional variation, indicating confounding between batch levels and shared latent scores. In contrast, residuals after MM-ComBat and MM-CovBat form more homogeneous clusters in the first two components, with MM-CovBat performing better in higher-order components.

#### Biological Covariance Pattern Simulation

To further investigate the whitening concern under different cross-metric covariance structures, we compared MM-ComBat (baseline covariance), MM-ComBat (target covariance), and SM-ComBat across all four scenarios. Figure 9 summarizes results across scenarios varying in the relative strength of biological and batch-related covariance.

**Figure 9.**
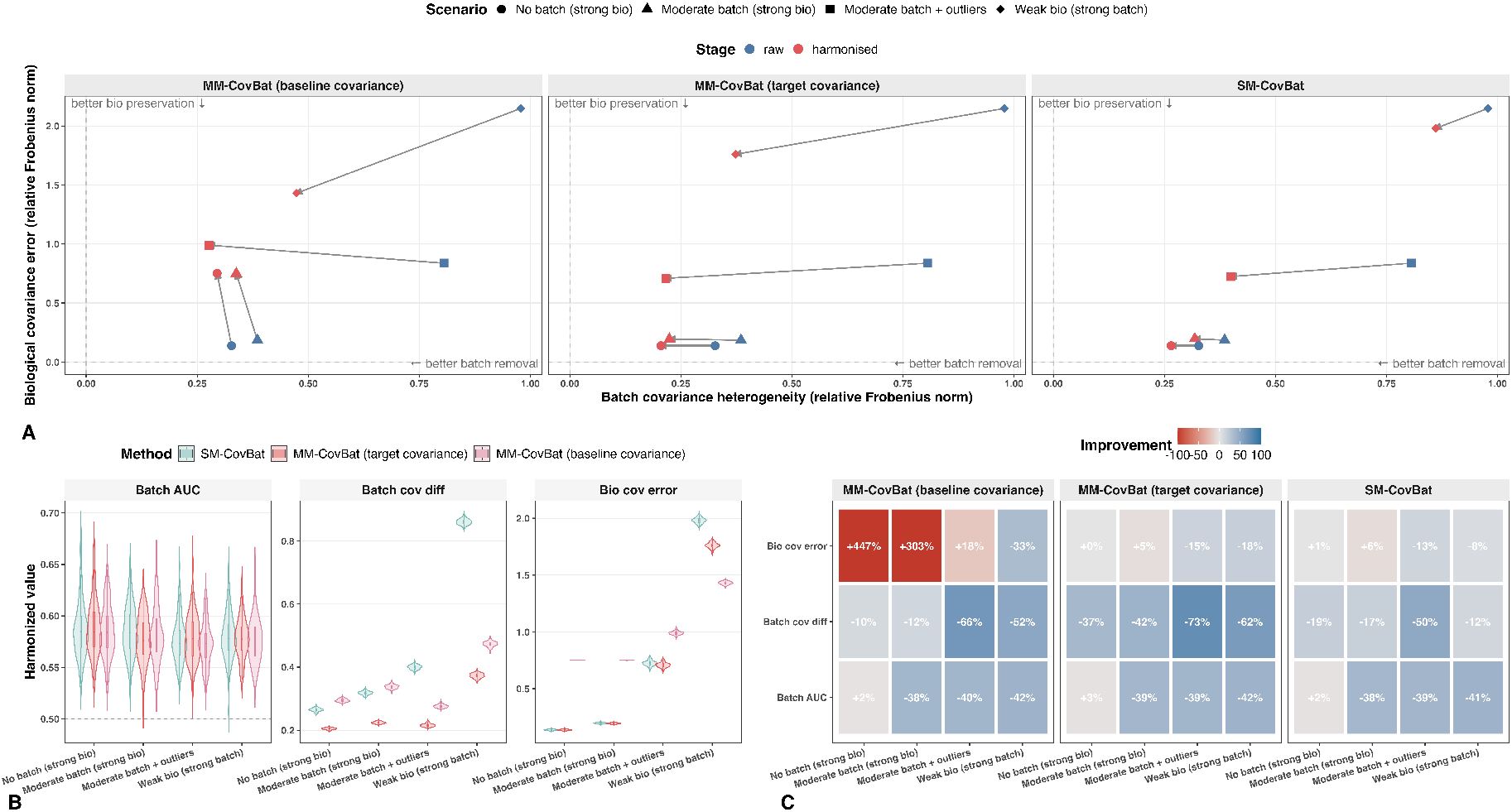
Harmonization performance in the biological-covariance simulation across four scenarios varying in the relative strength of biological and batch-related covariance. Results are averaged over 100 simulation replicates. (A) Trade-off between batch covariance heterogeneity and biological covariance error before and after harmonization. Lower values indicate better batch removal and better preservation of biological covariance. (B) Scenario-specific distributions of Batch AUC, batch covariance heterogeneity, and biological covariance error after harmonization. Lower Batch AUC indicates weaker residual batch detectability. (C) Mean percent change relative to the raw data for each method and scenario. Negative values indicate improvement for Batch AUC, batch covariance heterogeneity, and biological covariance error, whereas positive values indicate deterioration. Overall, MM-ComBat with target covariance provided the most consistent balance between reducing covariance-level batch effects and preserving biological covariance. In contrast, MM-ComBat with baseline covariance tended to over-correct when biological covariance was strong, but performed best in the scenario with strong batch structure and weak biological signal.

Figures 9B-C show that MM-ComBat (baseline covariance) tended to over-correct batch effects when strong biological patterns were present in the residual cross-metric covariance. In contrast, MM-ComBat (target covariance) kept this inflation well controlled. The exception was the strong-batch, weak-biology scenario, where MM-ComBat (baseline covariance) performed best. For batch covariance heterogeneity, MM-ComBat (target covariance) performed best across all scenarios, whereas MM-ComBat (baseline covariance) outperformed SM-ComBat in settings with severe outliers or strong covariance-level batch effects. Batch AUC was similar across the three methods, suggesting comparable removal of global batch signals.

The primary distinction between the two MM-ComBat formulations lay in the trade-off between batch removal and preservation of biologically meaningful covariance. Figure 9A shows that, in the scenarios characterized by strong biological covariance with either no batch covariance or moderate batch co-variance, MM-ComBat (baseline covariance) provided little additional gain in reducing batch effects in the cross-metric covariance, but substantially increased biological covariance error. In contrast, MM-ComBat (target covariance) removed more batch variation while keeping biological covariance error at a similar level, indicating better preservation of potential biological covariance patterns. SM-ComBat also kept biological covariance error well controlled, but achieved less batch removal than MM-ComBat (target covariance).

We observed a similar pattern in the scenario with severe outliers, though the inflation in biological covariance error under MM-ComBat (baseline covariance) was somewhat smaller than in the first two scenarios. However, in the scenario with strong batch effects and weak biological covariance, MM-ComBat (baseline covariance) performed best in terms of both batch removal and preservation of biological signal.

Overall, these results support using MM-ComBat (target covariance) when strong biological patterns remain in the cross-metric covariance, as the whitening concern for the baseline formulation is real and consequential in those settings. When the residual covariance is dominated by batch structure and biological covariance is weak, MM-ComBat (baseline covariance) is more appropriate. The attenuated over-correction under severe outliers may reflect outlier-driven distortion of the biological covariance itself, reducing the signal available to be inadvertently whitened. These results also bear directly on the question of how increasing marginal variances affect the whitening concern. Because the covariance-mapping step operates on covariance rather than correlation alone, larger marginal variances amplify covariance magnitudes even when the underlying correlation structure is unchanged. Whitening therefore removes proportionally more covariance mass as marginal variances grow, making the baseline formulation increasingly aggressive in settings where residual covariance is biologically driven. The target-covariance formulation mitigates this by remapping toward a shared structure rather than a structureless reference, and the biological covariance simulation described above was designed precisely to assess this regime.

To investigate the whitening concern for MM-CovBat, we compared MM-CovBat (baseline covariance), MM-CovBat (target covariance), and SM-CovBat across all four scenarios. The overall pattern was similar to that observed for MM-ComBat (Figure S7). In particular, MM-CovBat (target covariance) substantially reduced biological covariance error relative to MM-CovBat (baseline covariance), though slight inflation persisted when biological covariance was strong and batch covariance was weak or absent. Overall, MM-CovBat (baseline covariance) appeared more appropriate when the residual covariance was dominated by strong batch structure, whereas MM-CovBat (target covariance) was preferable when meaningful biological covariance structure was present.

## 5 Discussion

Standard harmonization approaches treat imaging metrics independently, leaving residual batch effects in cross-metric covariance that can distort downstream multivariate analyses. The methods developed here address this limitation directly. MM-ComBat jointly harmonizes correlated metrics to adjust for batch effects in means, variances, and covariances, while MM-CovBat extends this framework through latent-space harmonization to remove residual covariance-related batch effects. A target-covariance formulation complements the baseline approach by preserving shared biological covariance structure when that structure is strong, thereby reducing the risk of over-correction.

The A2CPS data confirmed that batch effects in multi-site neuroimaging extend beyond means and variances into cross-metric and cross-ROI covariance structure, consistent with prior evidence of persistent covariance discrepancies after standard site correction (Carmon et al. 2020). Under the baseline covariance formulation, MM-ComBat outperformed SM-ComBat in removing residual batch signal, particularly for noisy metrics where cross-metric borrowing stabilizes batch-effect estimation, while preserving more biological signal in the mean structure. MM-CovBat achieved the best overall performance across all batch-removal criteria, with EB providing robustness comparable to the more computationally expensive MCMC approach in this setting. Sensitivity analyses confirmed that the target-covariance formulation performed similarly to the baseline overall, with modest gains in covariance stabilization for outlier-prone metrics offset by slightly higher residual global batch signal.

In simulations with feature-wise batch effects and weak residual biological structure, both MM-ComBat variants under the baseline formulation reduced covariance-level batch effects while preserving biological variation in the mean structure, with the largest correlation recovery gains seen for cross-metric relationships under strong batch effects (Figure 5B). The FDR cost was modest and confined to weak signals (Figure 6).

Notably, although MM-ComBat assumes that batch effects are independent across features, cross-feature correlation recovery still improved. This likely reflects an indirect benefit of borrowing information across metrics, which stabilizes feature-wise batch-effect estimates and reduces spurious site-induced correlations.

When comparing the two estimation variants, the MCMC approach outperformed the EB variant in preventing within-metric correlation distortion under mild batch effects with small sample sizes. It also performed better in recovering cross-metric correlations under strong batch effects when priors were correctly specified. Notably, the MCMC variant retained strong performance even with misspecified priors under mild batch effects, likely because the likelihood from relatively clean data dominated and the Bayesian framework more effectively captured posterior uncertainty. Under more severe batch effects, the EB variant was more robust, whereas the MCMC approach became less stable and occasionally failed due to prior misspecification (Supplementary Figure S3). These failures were largely attributable to using weakly informative priors to model strongly structured covariance or mixtures of heterogeneous covariance regimes across features.

In the MCMC model, we used a separation strategy with half-*t* priors on marginal scales and an LKJ prior on the correlation matrix, yielding a flexible yet stable prior for the covariance matrix. When batch covariances were generated from an IW distribution, which is known to produce relatively unstructured covariance, our chosen priors aligned well with the true structure, substantially improving covariance recovery through multivariate pooling. In contrast, when covariances arose from structured families such as autoregressive (AR) or compound symmetry (CS), the exchangeable LKJ prior misrepresented the underlying patterns and over-shrank correlations toward independence (Supplementary Figure S6).

A similar issue arose when covariances were drawn from a mixture of heterogeneous covariance structures across features. The LKJ prior imposed an implicit exchangeability assumption that encouraged overly similar correlation patterns across features and attenuated feature-specific structure. Under such prior–likelihood mismatches, particularly in settings with limited sample size or strong batch effects, posterior inference became strongly regularized by the prior, leading to under-recovery of covariance structure or inflated Type I error rates. To address this limitation and enhance flexibility across diverse covariance structures, future extensions of MM-ComBat should incorporate more flexible covariance priors capable of adapting to both unstructured and structured dependence patterns, as well as mixtures of heterogeneous covariance structures across features.

Beyond feature-wise batch effects, MM-CovBat showed superior performance under latent-space batch effects in removing batch effects in feature correlations both within and across metrics, with corresponding gains in biological signal separation (Figure 8A). MM-ComBat remained robust in mitigating batch effects in feature correlations, even when the assumed batch-effect distribution was violated. This robustness likely reflects its multivariate modeling strategy, which borrows information across metrics and facilitates cross-feature correlation recovery even when batch effects interact with biological signals in some metrics. In contrast, SM-CovBat, treating metrics independently, struggled to disentangle batch from biological signals under strong confounding (Figure 8B), highlighting the advantage of joint modeling.

When biological covariance is strong and batch effects are moderate, the baseline formulation can whiten biologically meaningful cross-metric correlations along with batch effects. The target-covariance formulation addresses this by remapping the adjusted covariance toward a shared estimated covariance structure, thereby preserving common biological variation across batches while still reducing batch effects. In simulations with structured biological covariance, the target formulation largely retained the batch-removal ability of the baseline formulation while substantially reducing distortion of the biological covariance structure (Figure 9). The choice between formulations therefore depends on the relative strength of biological and batch-related covariance in the residuals and should be guided by diagnostic assessment before harmonization. A practical first step is to examine cross-metric correlation matrices both at the level of individual features and after averaging across features, as illustrated in Figure S11. Stable off-diagonal patterns present within batches and in the pooled sample suggest strong shared biological structure, favoring the target-covariance formulation, whereas a more diagonal average matrix with greater between-batch heterogeneity in feature-specific correlations or stronger pooled than within-batch correlations suggests predominantly batch-driven covariance, favoring the baseline. For a more formal assessment, projecting harmonized residuals onto the eigenvectors of the target covariance and testing for residual site differences can confirm whether the estimated target retains batch-related signal and whether falling back to the baseline formulation is warranted. The trade-off is that the remapping step may partially reintroduce batch effects if the target covariance estimate is contaminated. The baseline formulation remains preferable when batch covariance is strong, while the target formulation is more advantageous when biological covariance is pronounced. Resolving this tradeoff fully would require either an external reference dataset known to be free of batch effects (from which a purely biological covariance target could be estimated independently) or a model that jointly parameterizes batch-related and biological covariance as distinct components. Neither is available in the typical multi-site setting without strong additional assumptions, and the separation of batch from biological covariance is therefore fundamentally constrained by the available data structure. The target-covariance formulation is accordingly best understood as a practical approximation that exploits shared cross-batch structure as a proxy for non-batch covariance, rather than a complete solution to this identifiability challenge.

It is worth noting that the whitening concern addressed by the target-covariance formulation is not unique to MM-ComBat and MM-CovBat. Instead, it is an inherited limitation of any ComBat-family method that includes a covariance harmonization step, including the original CovBat, which harmonizes within-site PC score variances toward a pooled value and faces the same issue when between-site covariance differences are small or biologically driven. The target-covariance formulation introduced here addresses this broader limitation in principle for any such method, provided that a shared biological target can be estimated from the pooled data. To our knowledge, no prior work in this framework has proposed a practical mechanism for preserving biologically meaningful covariance during harmonization, and we therefore view this as a contribution that extends beyond the specific multivariate extensions described in this paper.

Together, these findings indicate that the choice of harmonization method should be guided by both diagnostic evidence and underlying data structure. We therefore propose a practical workflow for selecting among single-metric and multi-metric harmonization strategies (Figure 10). The workflow summarizes how feature-level, global, and covariance-related diagnostics, together with the feasibility of joint multi-metric analysis, inform the choice among ComBat-family methods, CovBat, MM-ComBat, and MM-CovBat.

**Figure 10.**
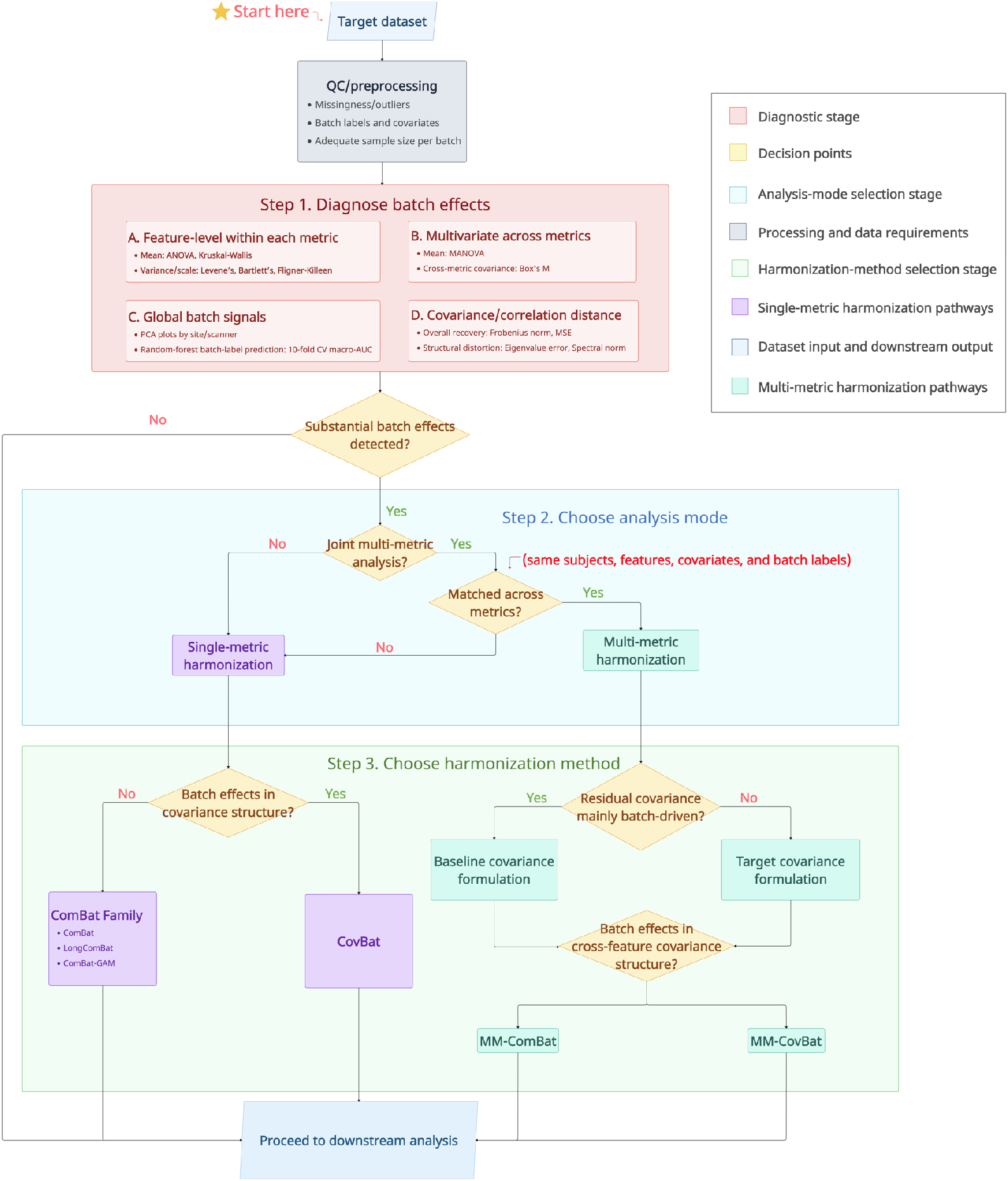
A practical workflow for selecting a harmonization strategy. Step 1 consists of preprocessing and diagnostic assessment of batch effects at the feature, global, and covariance levels. Step 2 determines whether the data support single-metric or joint multi-metric harmonization. Step 3 identifies the appropriate harmonization method. For single-metric analysis, covariance-structure batch effects guide the choice between the ComBat family and CovBat. For multi-metric analysis, the relative contribution of batch-driven versus biologically driven covariance guides the choice between the baseline and target covariance formulations, and residual cross-feature covariance batch effects guide the choice between MM-ComBat and MM-CovBat. If no substantial batch effects are detected, the data can proceed directly to downstream analysis.

Despite this practical guidance, several limitations should be considered when applying these methods. As in the original ComBat framework, our methods require pre-specification of covariates whose effects are to be preserved as biologically meaningful signal. Prior work has shown that ComBat-based harmonization cannot preserve the effects of covariates that are not explicitly included in the model, particularly when those covariates are confounded with site (Bayer et al. 2022). Consequently, omitting a relevant covariate may result in its associated variation being inadvertently removed along with batch effects. We compared the performance of single-metric and multi-metric harmonization frameworks in preserving the effect of an omitted covariate in the mean structure using both the A2CPS dataset, which provides a realistic setting with substantial confounding, and simulations in which the omitted covariate was designed to have effects that were not confounded with batch. We found that both MM-ComBat and MM-CovBat under the baseline covariance formulation tended to remove unprotected biological signals more aggressively when strong confounding was present, as observed in the A2CPS data (Supplementary Figure S5A,B). In contrast, the single-metric framework showed more consistent preservation of unprotected biological variation, likely because it shares less information across features and no information across metrics. However, when confounding was weak or absent, the two multi-metric frameworks performed better at preserving unprotected biological signals by yielding more accurate batch-effect estimates, with MM-CovBat performing slightly better overall (Supplementary Figure S5C). This concern is particularly relevant when demographic variables are not well balanced across sites, since strong confounding between covariates and batch makes it more difficult for any ComBat-based method to preserve biologically meaningful effects while removing site-related variation. These results therefore highlight the importance of thorough exploratory analyses and batch-effect diagnostics before applying multivariate harmonization, so that covariates to be preserved can be specified appropriately and concerns about unintentionally removing important biological signals can be minimized.

Furthermore, the current framework handles only one batch variable at a time, posing challenges for datasets with hierarchical batch structures, such as nested scanner parameters or software updates. Some studies have combined batch factors pairwise (Beer et al. 2020), but this treats each batch level as independent and ignores potential information sharing (e.g., across scanners from the same vendor). A generalized ComBat method that adjusts batch variables sequentially has shown promise (Horng et al. 2022), yet sequential adjustment may disrupt dependencies among batch variables and amplify noise due to accumulated estimation uncertainty. Future work should aim to incorporate hierarchical structures into harmonization models. In addition, because our real-data application focused on cortical metrics, further validation on other highly correlated neuroimaging modalities, such as diffusion-derived measures, will be important for assessing the generalizability of the proposed multivariate framework beyond this setting.

The current multi-metric longitudinal ComBat should be viewed as a working solution. It relies on the strong assumption that a mixed-effects model adequately captures within-subject dependence, so that the resulting residuals can be treated as independent observations for batch estimation. In high-dimensional settings this assumption is likely to be violated, and a more rigorous multivariate LongComBat with explicit longitudinal covariance modeling remains an important direction for future work.

A further practical consideration concerns the standardization step preceding batch-effect estimation. The current implementation offers a default standardization based on pooled residual variance and a robust alternative using median centering and Tukey’s biweight midvariance. The latter is intended primarily to reduce sensitivity to outliers and improve stability in heavy-tailed settings. Because both pool across batches, the scale estimate may retain some batch-related heterogeneity. In most settings, this is unlikely to materially affect harmonization performance. However, when batch-effect magnitudes differ substantially across metrics, it may modestly bias metric-specific calibration. Batch-aware scaling strategies that more explicitly separate residual scale from batch variation are a natural target for future development.

Finally, pooling across metrics may inflate the FDR if non-null signal in some metrics propagates to weaker or null effects in others. Careful selection of metrics for joint harmonization is therefore important. The current MM-ComBat (MCMC) implementation uses a two-stage framework in which fixed effects are first removed and batch effects are then estimated. Although computationally efficient, this design does not fully exploit the Bayesian framework’s ability to jointly estimate fixed and batch effects. Misspecification of the regression model may bias fixed-effect estimates and, in turn, affect batch correction. Future work should explore fully Bayesian models that simultaneously estimate fixed and batch effects and compare their performance with the proposed two-stage approach.

## Data and Code Availability

Data were provided by the A2CPS Consortium funded by the National Institutes of Health (NIH) Common Fund, which is managed by the Office of the Director (OD)/ Office of Strategic Coordination (OSC). Consortium components and their associated funding sources include Clinical Coordinating Center (U24NS112873), Data Integration and Resource Center (U54DA049110), Omics Data Generation Centers (U54DA049116, U54DA049115, U54DA049113), Multi-site Clinical Center 1 (MCC1) (UM1NS112874), and Multi-site Clinical Center 2 (MCC2) (UM1NS118922). The data release used in this analysis is available for download from the NIMH Data Archive (NDA).

We integrated SM-ComBat, SM-CovBat, MM-ComBat, and MM-CovBat into a unified R package, `MultiComBat`, which also includes a suite of commonly used tools for diagnosing batch effects. The design of `MultiComBat` builds upon the univariate ComBat-family framework as implemented in existing R software `ComBatFamily` (Chen and Gardner 2024), and extends this framework to the multivariate setting while providing a unified interface and additional diagnostic functionality. The package is available on GitHub at https://github.com/Zheng206/MultiComBat.

## Author Contributions

**Zheng Ren**: Conceptualization, Methodology, Software, Validation, Formal Analysis, Investigation, Resources, Data Curation, Writing - Original Draft, Writing - Review & Editing, Visualization. **Patrick Sadil**: Data Curation, Writing - Original Draft, Writing - Review & Editing, Visualization. **Martin A. Lindquist**: Conceptualization, Methodology, Writing - Original Draft, Writing - Review & Editing, Supervision, Project Administration, Funding Acquisition.

## Ethics Statement

Informed consent was obtained from all A2CPS participants.

## Acknowledgements

We thank the developers of the `ComBatFamily` R package for providing an open implementation of the ComBat-family framework that informed the development of `MultiComBat`.

## Appendix A Derivation for Multivariate EB Batch Adjustment

### A.1 Framework Specification

In the multivariate ComBat framework, let ***Z***_*ijv*_ ∈ ℝ^*M*^ denote the rescaled residual vector of *M* metrics for feature *v* from batch *i* and subject *j*, after removing fixed effects. The model assumes the following:

1. **Data Likelihood:** The observed data for each feature *v* in batch *i* follow a multivariate normal distribution with mean ***γ***_*iv*_ and covariance **Σ**_*iv*_:

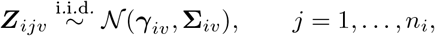

where ***γ***_*iv*_ ∈ ℝ^*M*^ is the batch-specific location effect, and **Σ**_*iv*_ ∈ ℝ^*M* ×*M*^ is a symmetric positive-definite covariance matrix that captures the batch-specific dispersion across the *M* metrics for feature *v* in batch *i*, shared across subjects *j. n*_*i*_ is the number of samples in batch *i*.
2. **Prior for *γ***_*iv*_ : The mean ***γ***_*iv*_ for each feature *v* in batch *i* is modeled with a multivariate normal prior:

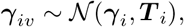

where ***γ***_*i*_ ∈ ℝ^*M*^ is a batch-level mean and ***T*** _*i*_ ∈ ℝ^*M* ×*M*^ is a covariance matrix that governs how batch means ***γ***_*iv*_ vary *across features v* within batch *i*.
3. **Prior for Σ**_*iv*_: The covariance matrix **Σ**_*iv*_ for each feature in batch *i* follows an Inverse-Wishart distribution:

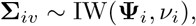

where **Ψ**_*i*_ is the scale matrix and *ν*_*i*_ is the degrees of freedom.

### A.2 Estimating Hyperparameters

We estimate the prior hyperparameters (***γ***_*i*_, ***T*** _*i*_, **Ψ**_*i*_, *ν*_*i*_), and initialize ***γ***_*iv*_ and **Σ**_*iv*_, using the method of moments.

#### Sample-specific Estimates

The sample-specific mean 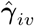 and variance 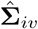 estimates are calculated as follows:

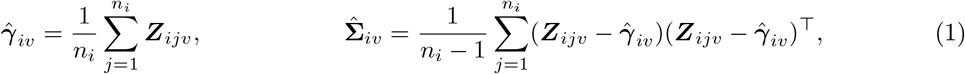

#### Multivariate Normal Prior Hyperparameter Estimates

The estimates for the hyperparameters of the normal prior are computed as follows:

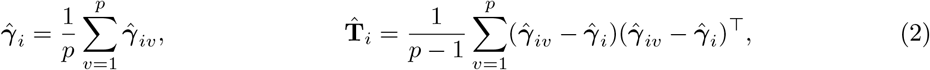

where *p* is the total number of features.

#### Inverse-Wishart Prior Hyperparameter Estimates

Moment matching for the Inverse–Wishart prior is a bit more complicated due to the second moments. To simplify, we match only the first moment and the diagonal variances. The estimates for the hyperparameters of the Inverse-Wishart prior are computed as follows:

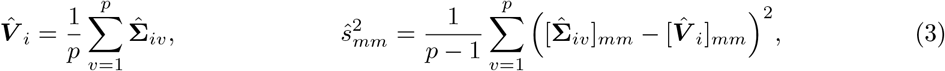

where [·]_*mm*_ denotes the (*m, m*) element and *m* = 1, …, *M*.

The sample moments 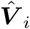 and 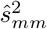 are then used to match the theoretical first moment and diagonal second moment of the Inverse-Wishart distribution. We average across diagonals to obtain a scalar degrees-of-freedom estimate:

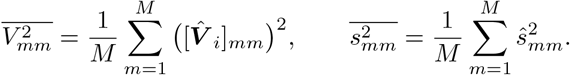

By solving the moment equations, we obtain

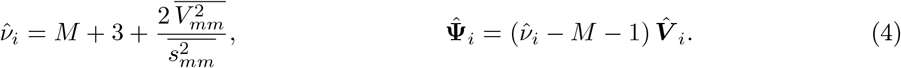

### A.3 Obtaining EB Batch Effects

Once the method-of-moments (MoM) hyperparameters are obtained, we apply an empirical Bayes procedure to pool information across features within each batch. This shrinks the feature-specific batch parameters toward the batch-level mean. Specifically, under the priors ***γ***_*iv*_ ~ 𝒩 (***γ***_*i*_, ***T*** _*i*_) and **Σ**_*iv*_ ~ IW(**Ψ**_*i*_, *ν*_*i*_), the conditional posteriors are available in closed form.

**Posterior Distribution for *γ***_*iv*_

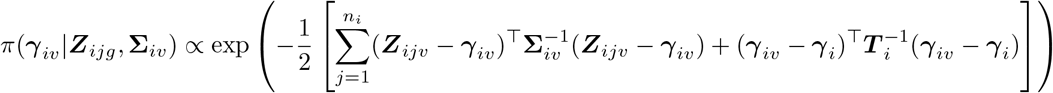

Therefore, the posterior mean of ***γ***_*iv*_ can be expressed as:

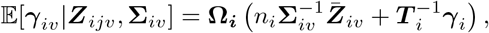

where:

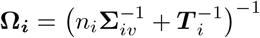

and

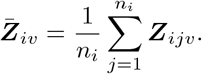

Given 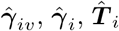, and 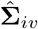 as defined above, the posterior mean can be estimated as:

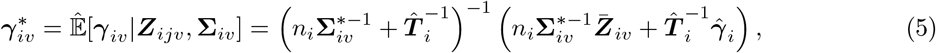

**Posterior Distribution for Σ**_*iv*_

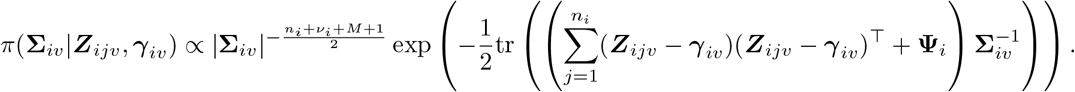

The posterior mean for **Σ**_*iv*_ is then:

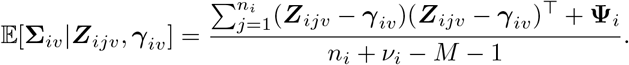

Given 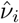 and 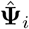 as defined above, the posterior mean can be estimated as:

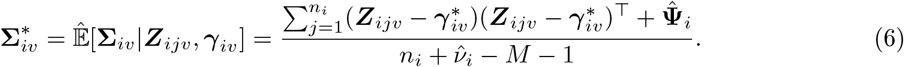

## B Derivation of G Space via the Rayleigh-Ritz Theorem

We consider

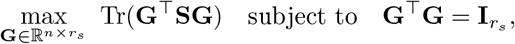

where 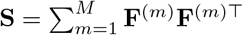 is symmetric and positive semidefinite.

Let **S** = **UΛU**^⊤^ be its eigendecomposition, with **Λ** = diag(*λ*_1_, …, *λ*_*n*_) and *λ*_1_ ≥ · · · ≥*λ*_*n*_ ≥0. Substituting gives

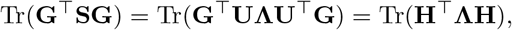

where **H** = **U**^⊤^**G** and 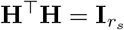.

By the Rayleigh-Ritz theorem, this quantity is maximized when the columns of **H** select the top *r*_*s*_ eigenvectors of **Λ**, yielding

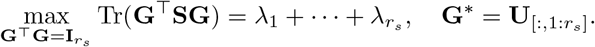

Hence, the maximizer **G**^∗^ consists of the top *r*_*s*_ eigenvectors of

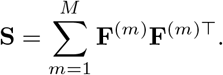

## C Robust Estimation of Shared Covariance

Under the target-covariance formulation, we assume that the residual covariance reflects a mixture of shared biological structure, batch-related variation, and random noise. Let the residual vector for subject *j* in batch *i* and feature *v* be

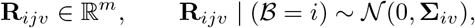

where ℬ denotes batch membership. At the covariance level, we decompose the batch-specific covariance as

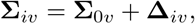

where **Σ**_0*v*_ denotes the shared covariance target and **Δ**_*iv*_ represents the batch-specific deviation. To make this decomposition identifiable, we assume the weighted centering condition

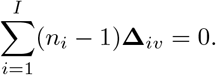

If *P* (ℬ = *i*) = *π*_*i*_, then by the law of total covariance,

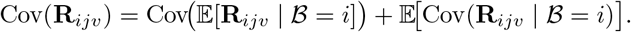

Because 𝔼[**R**_*ijv*_ | ℬ = *i*] = 0, this simplifies to

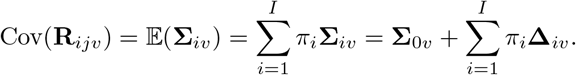

Thus, if the batch-specific deviations are centered in the population so that

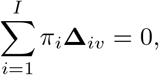

then

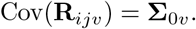

That is, **Σ**_0*v*_ can be interpreted as the population-average residual covariance across batches. Motivated by this decomposition, a natural estimator of **Σ**_0*v*_ is the degrees-of-freedom-weighted average

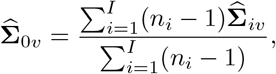

where 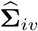 is the sample covariance matrix in batch *i*. Since

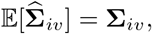

we have

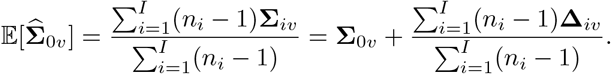

Therefore, under the weighted centering condition

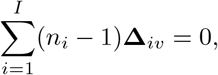

the pooled estimator is unbiased:

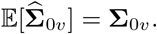

This provides the idealized justification for using a pooled covariance matrix as an estimator of the shared covariance target. In practice, however, covariance estimates may be unstable, particularly when batch sizes are small or the data are noisy. To improve stability and reduce the influence of aberrant covariance estimates, we replace the sample covariance matrices 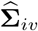 with EB-stabilized estimates 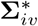 and further apply robust batch-specific weights. The resulting estimator is no longer exactly unbiased in the classical sense, but it is less sensitive to outlying batch-specific covariance patterns and can provide a more stable estimate of the shared covariance target in finite samples.

## Supplementary Materials

### S1 Bayesian Model and MCMC Implementation

The multivariate Bayesian model was fit using Markov Chain Monte Carlo (MCMC) sampling implemented in Stan. We ran three independent chains in parallel, with an adaptive Hamiltonian Monte Carlo sampler using a target acceptance rate of 0.98. Batch-level intercepts were assigned standard normal priors. Feature- and batch-specific mean vectors were modeled with normal priors centered on the corresponding batch intercepts. Standard deviation parameters were assigned half-Student-*t* priors with degrees of freedom drawn from a Gamma(2, 0.1) distribution and scale parameters drawn from a Cauchy(0, 2.5) distribution. Correlation structures were modeled using LKJ priors with shape parameter 1. A complete specification of prior distributions is provided in Supplementary Table S6. Convergence was assessed using standard diagnostics provided by Stan, including the potential scale reduction factor 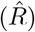 and effective sample size.

### S2 Sensitivity Analysis

In this section, we conduct a sensitivity analysis to evaluate the potential whitening concern in a real neuroimaging application and to assess how the target-covariance formulation influences harmonization results. Specifically, we applied the target-covariance versions of MM-ComBat and MM-CovBat to the A2CPS dataset and evaluated them using the same diagnostic framework as in the main analysis, including feature-wise batch removal, preservation of biological signal in the mean structure, residual global batch signal within each metric, and batch effects in correlation structure. Overall, the conclusions were broadly consistent with those obtained under the baseline covariance formulation. In particular, the whitening concern did not materially alter conclusions regarding batch removal in the mean structure or preservation of biological effects modeled through the prespecified regression design. Nevertheless, several differences emerged, highlighting both the potential advantages and limitations of mapping residual covariance to an estimated common target structure.

As shown in Figure S8B, the target-covariance formulation performed nearly identically to the baseline formulation in removing feature-wise batch effects in cross-metric covariance, indicating that the whitening concern has little impact on this aspect of performance. For within-metric feature-wise batch effects (Figure S8A), the target-covariance formulation showed slight improvement in removing multiplicative batch effects for metrics with more severe outliers, such as FoldInd and CurvInd. For residual global batch signal (Figure S9B), MM-CovBat under the target-covariance formulation continued to perform best across metrics. In contrast, MM-ComBat under the target-covariance formulation appeared to retain some batch-related covariance structure for metrics with severe outliers, with AUC values around 0.75–0.80. A plausible explanation is that the estimated target covariance may contain both biologically meaningful shared structure and residual batch-related structure. Consequently, reintroducing this common covariance may partially restore unwanted batch variation. Even so, both MM-ComBat variants continued to outperform SM-ComBat and SM-CovBat, supporting the advantage of multi-metric harmonization over single-metric approaches for batch removal.

With respect to preservation of biological signal, we observed the same general pattern as under the baseline formulation: MM-ComBat and MM-CovBat tended to yield more statistically significant biological associations, with MM-CovBat showing the strongest overall performance (Figure S9A). Relative to the baseline covariance formulation, however, all multi-metric methods showed slightly weaker apparent signal recovery. This may indicate that the target-covariance mapping step reintroduces residual variation that is not purely biological, thereby modestly reducing power. Alternatively, it may suggest a reduction in false positives if the baseline formulation removes covariance structure too aggressively. Because these comparisons are based on downstream association results rather than direct ground truth, we interpret this pattern as modest attenuation in apparent signal recovery rather than definitive evidence of true biological signal loss.

Finally, the target-covariance formulation substantially improved removal of batch effects in within-metric feature correlation, particularly for MM-CovBat. Under the baseline covariance formulation, SM-CovBat showed the strongest performance across all four within-metric covariance criteria (Frobenius norm, MSE, spectral norm, and eigenvalue error). Under the target-covariance formulation, however, both MM-ComBat methods moved much closer to SM-CovBat across these criteria, while MM-CovBat outperformed SM-CovBat in Frobenius norm, MSE, and eigenvalue error (Figure S10A). These results suggest that the target-covariance formulation can improve harmonization of within-metric covariance structure, likely by preserving shared covariance patterns that would otherwise be overly shrunk or removed under the baseline whitening approach. By contrast, performance for cross-metric covariance harmonization remained largely unchanged between the two covariance formulations (Figure S10B).

In summary, the baseline and target-covariance formulations produced broadly similar results overall, especially for removal of batch effects in cross-metric covariance. The main advantage of the target-covariance formulation appears to be improved harmonization of within-metric feature covariance, potentially because it better preserves meaningful shared covariance structure. At the same time, if the estimated target covariance retains residual batch-related patterns, this approach may reintroduce some unwanted batch signal. Thus, under the current formulation, the target-covariance approach involves a trade-off between covariance preservation and complete batch removal, and this trade-off should be considered when interpreting results.

### S3 Supplementary Figure

**Figure S1.**
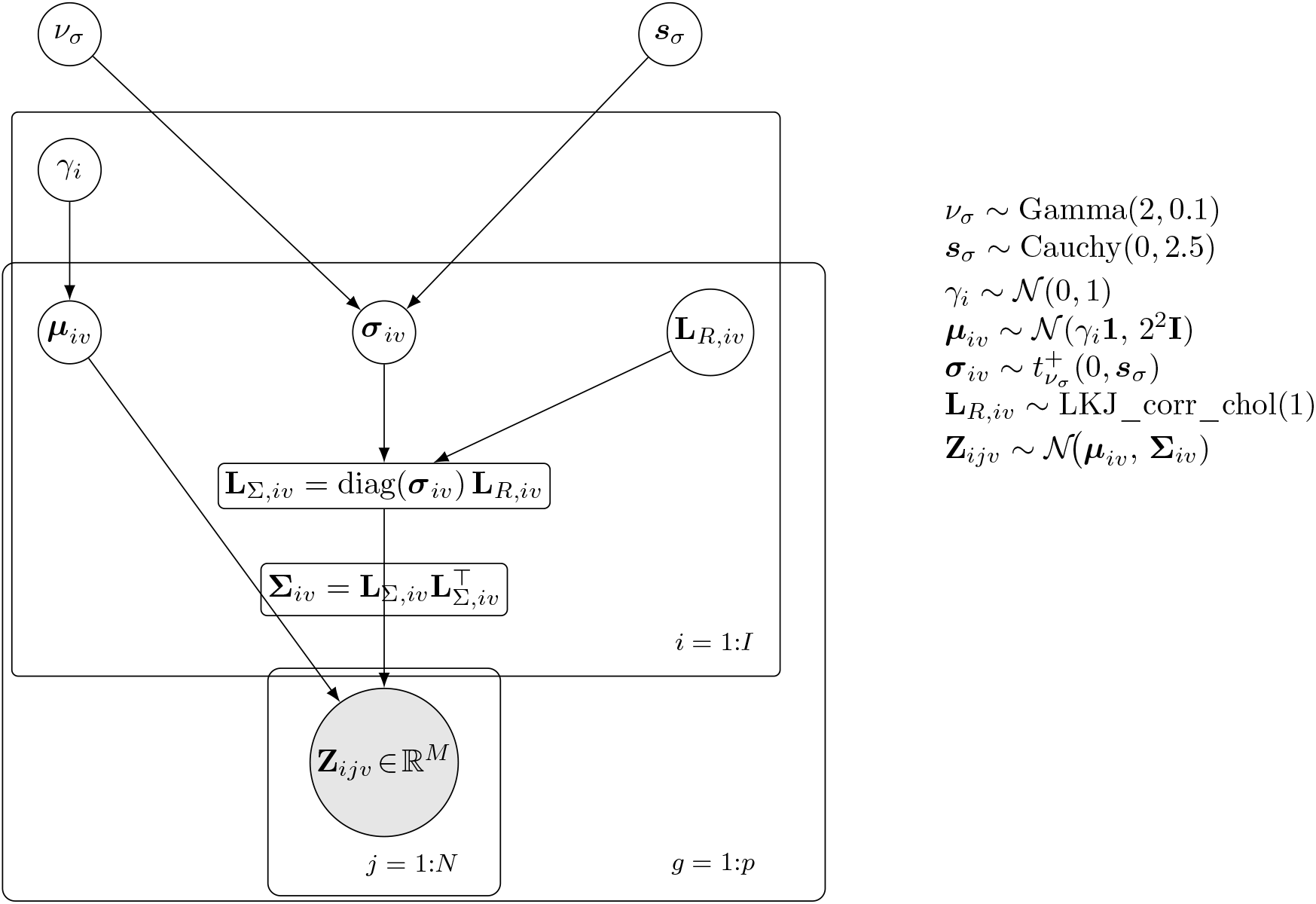
Plate diagram corresponding to the Stan model. Shaded = observed, white = latent, rounded rectangle = deterministic transform. Bold symbols are *M*-vectors.

**Figure S2.**
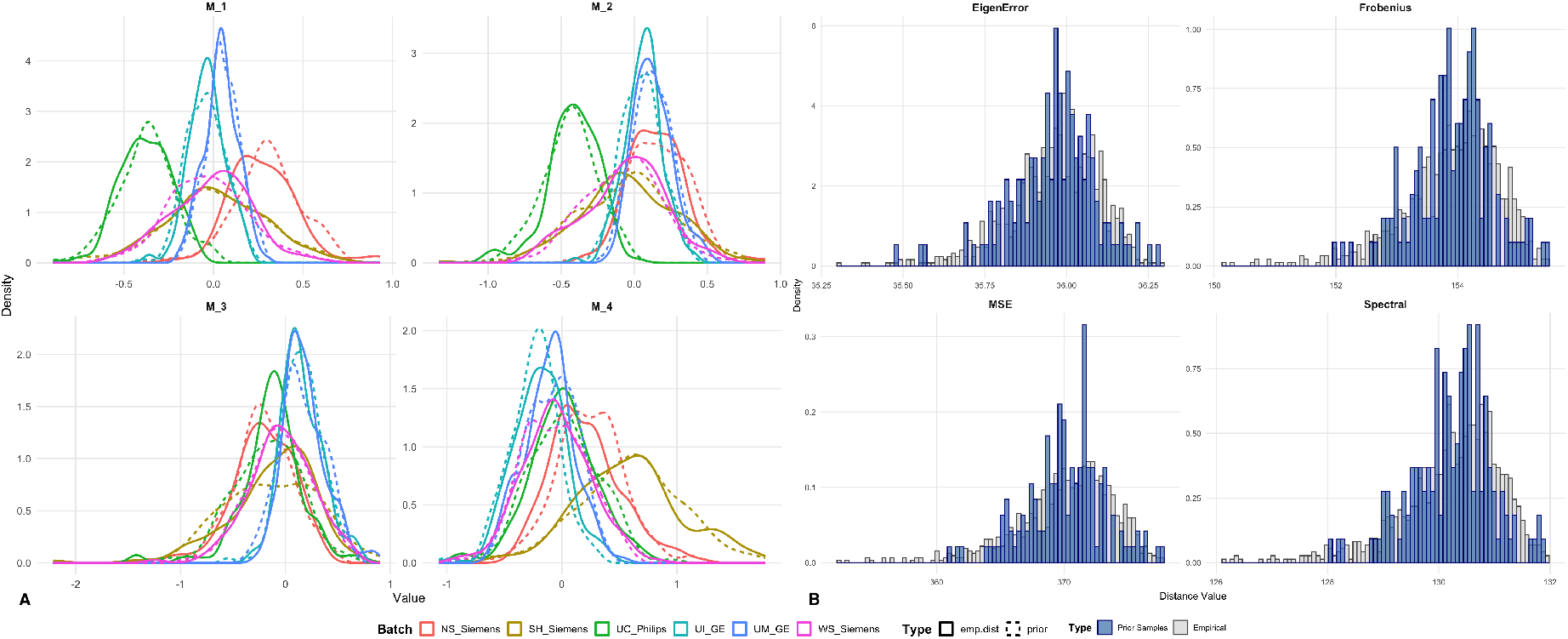
EB Assumption Check. (A) Comparison of the empirical distribution of estimated additive batch effects with the corresponding gamma prior. (B) Prior predictive check for the IW prior used for the multiplicative batch effects.

**Figure S3.**
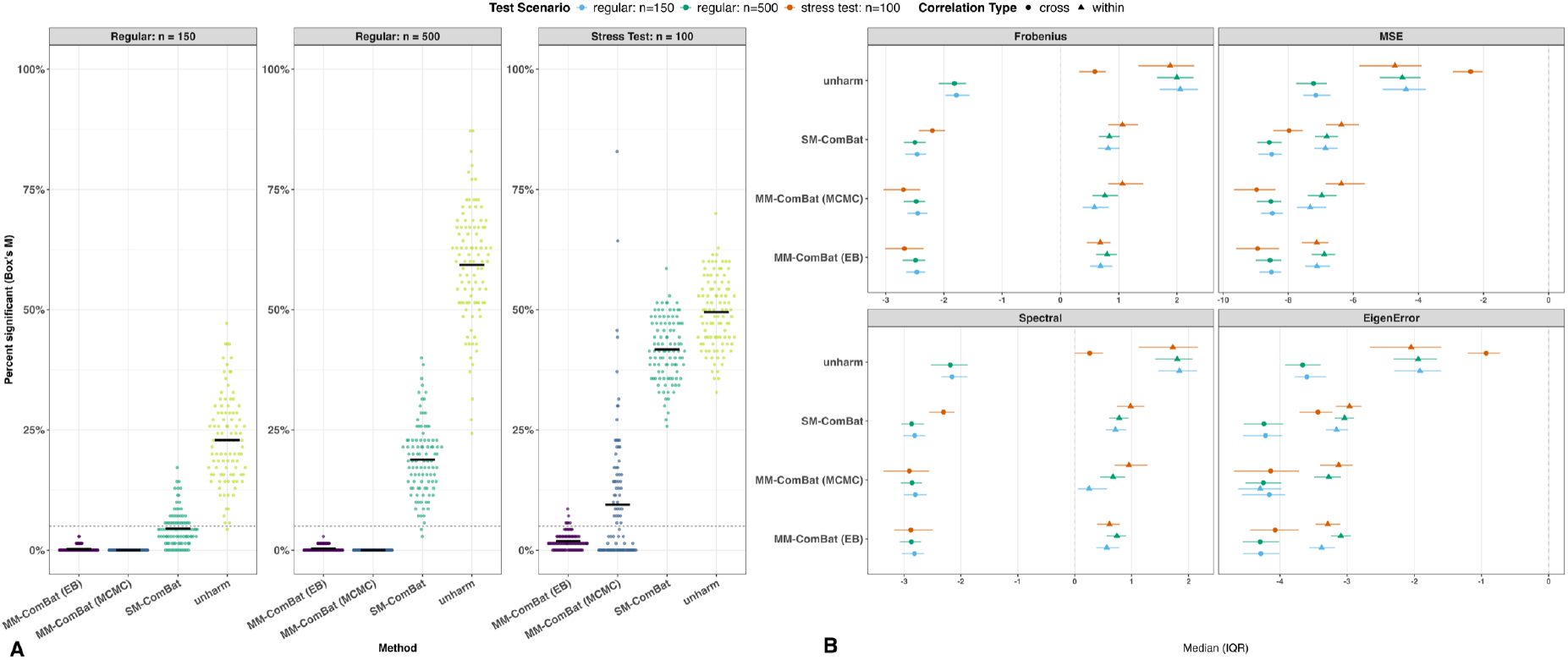
Batch Detection in Covariance and Correlation Recovery (Model-Misspecified; c.f., Figure 4). (A) Box’s M test across experimental conditions. Both MM-ComBat variants outperformed SM-ComBat, with the MCMC variant showing greater gains under mild batch effects, but becoming less stable under stress. (B) Correlation recovery. The MCMC approach performed similarly to the EB approach.

**Figure S4.**
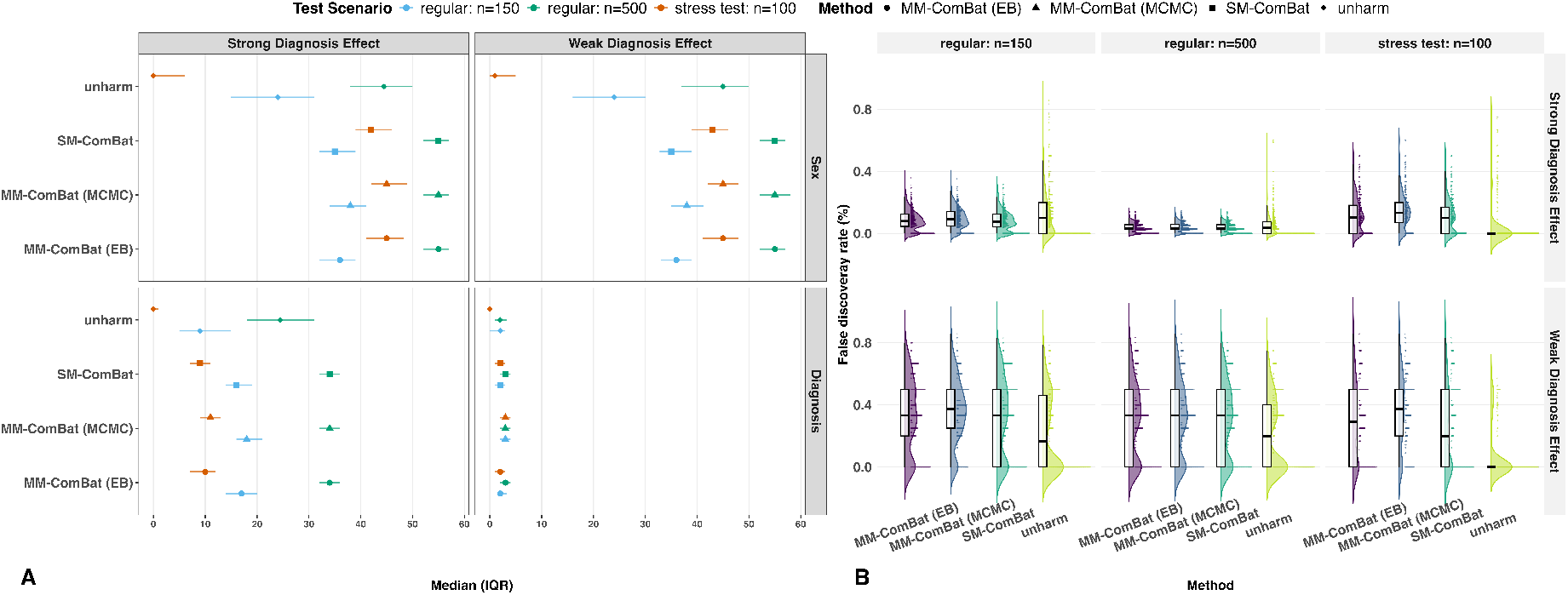
Fixed-effects Detection and Corresponding FDRs (Model-Misspecified; c.f., Figure 6). (A) Preservation of biological signals. Counts of features with significant sex and diagnosis effects across experimental conditions. MM-ComBat (MCMC) performed similarly to MM-ComBat (EB). (B) False discovery rate (FDR) based on known biomarkers. Both MM-ComBat variants performed comparably to SM-ComBat.

**Figure S5.**
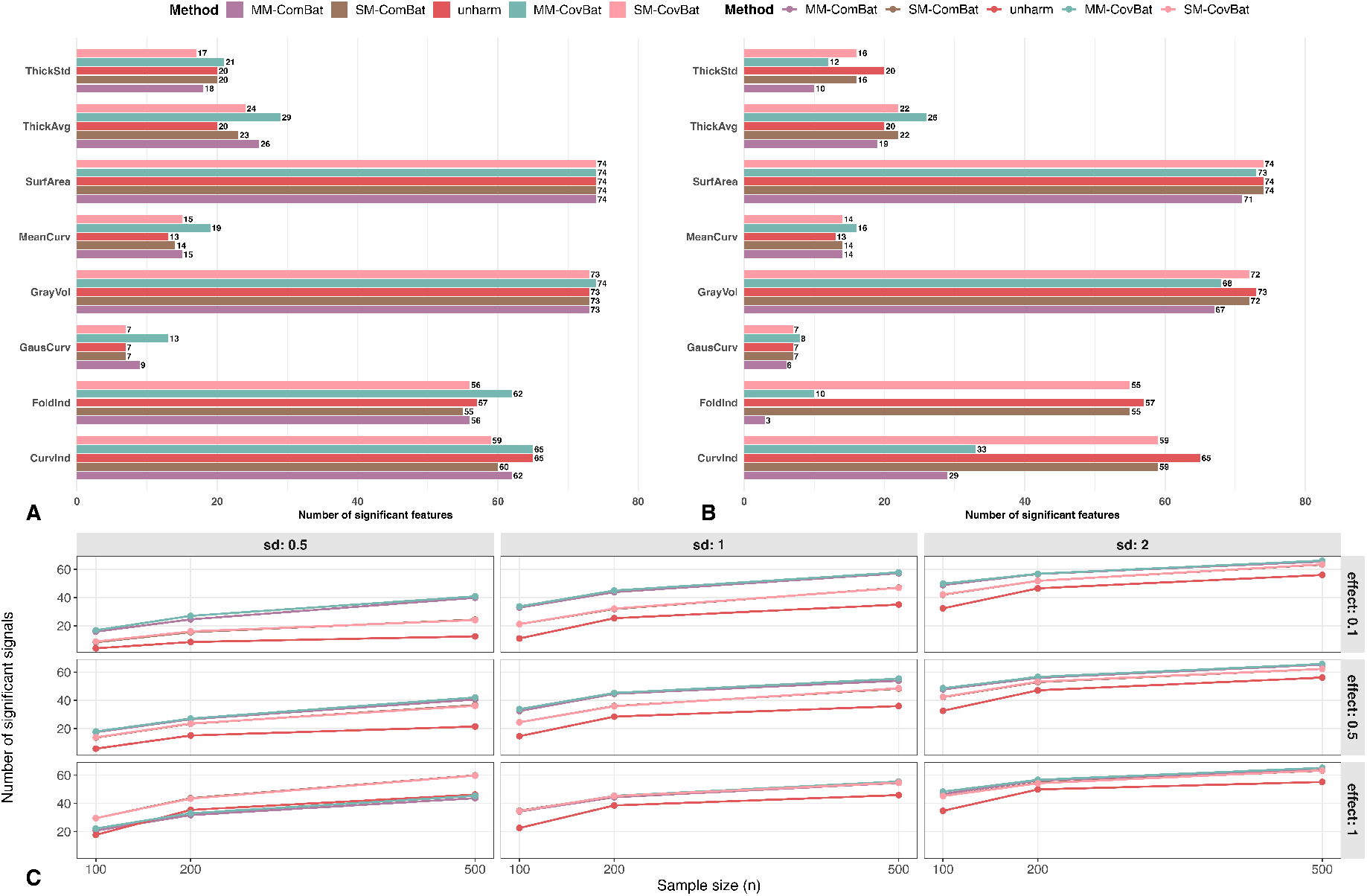
Biological signal preservation. (A) Preservation of sex effects in the A2CPS data when sex is included as a covariate, shown as the number of features with significant sex effects. (B) Preservation of sex effects in the A2CPS data when sex is not included as a covariate, shown as the number of features with significant sex effects. (C) Preservation of an omitted biological signal in simulations varying by effect size, signal standard deviation, and sample size.

**Figure S6.**
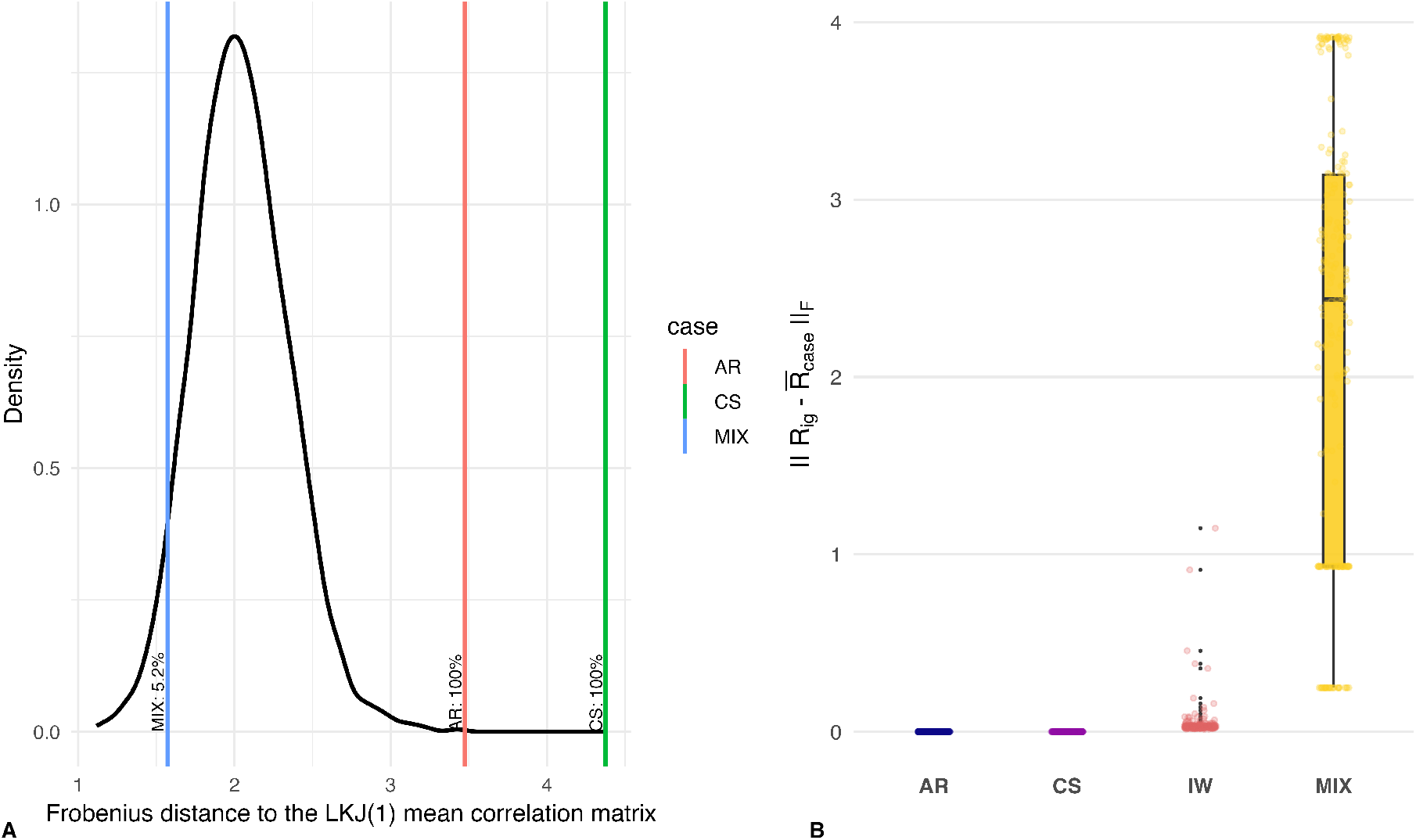
Prior-data alignment and feature-wise heterogeneity across covariance regimes. (A) Distribution of Frobenius distances between correlation matrices drawn from an LKJ(1) prior and the LKJ(1) mean correlation matrix (black density). Vertical lines indicate the Frobenius distance between the case-averaged simulated correlation matrix and the LKJ(1) mean for each data-generating regime (AR, CS, MIX). Correlations generated under the AR and CS regimes lie far outside the typical support of the LKJ(1) prior, indicating substantial prior–data mismatch. In contrast, the MIX regime exhibits a markedly smaller distance, which arises from averaging across heterogeneous, feature-specific covariance regimes rather than from genuine alignment with the LKJ prior. (B) Feature-wise deviations of individual correlation matrices from the case-average correlation. While the AR and CS regimes show minimal within-case variability, the MIX regime exhibits extreme feature-level heterogeneity, with large dispersion across features. Taken together, these patterns help explain why exchangeable priors such as the LKJ can obscure feature-specific structure and degrade performance under heterogeneous covariance regimes, particularly in data-limited or high-noise settings.

**Figure S7.**
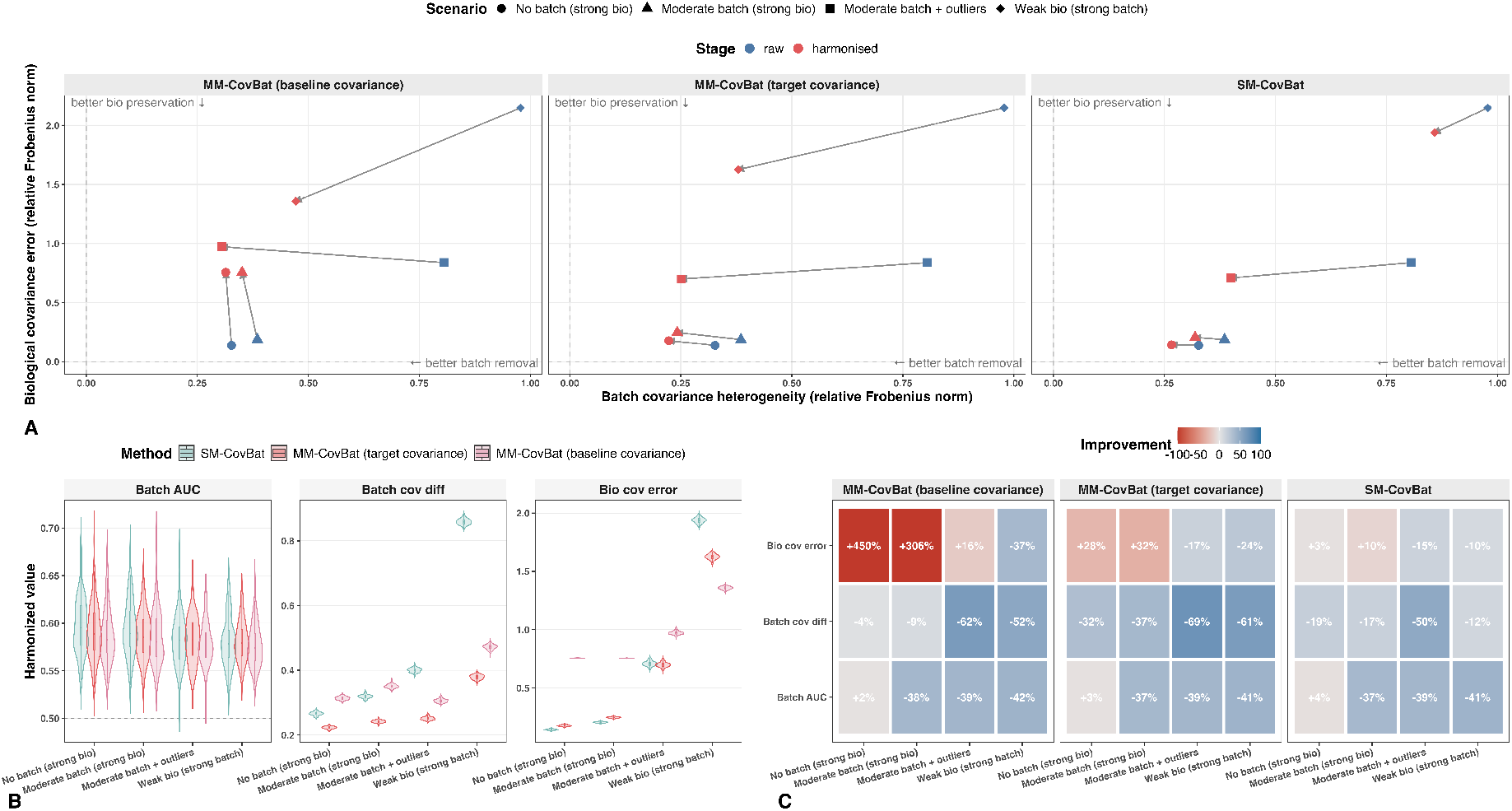
Harmonization performance in the biological-covariance simulation across four scenarios varying in the relative strength of biological and batch-related covariance for MM-CovBat. Overall, MM-CovBat with the target-covariance formulation provided the most consistent balance between reducing covariance-level batch effects and preserving biological covariance. Relative to the baseline-covariance formulation, it substantially reduced inflation in biological covariance error, although slight inflation remained when batch covariance was weak or absent. By contrast, the baseline-covariance formulation tended to overcorrect when biological covariance was strong, but performed best when residual covariance was dominated by strong batch structure and weak biological signal.

**Figure S8.**
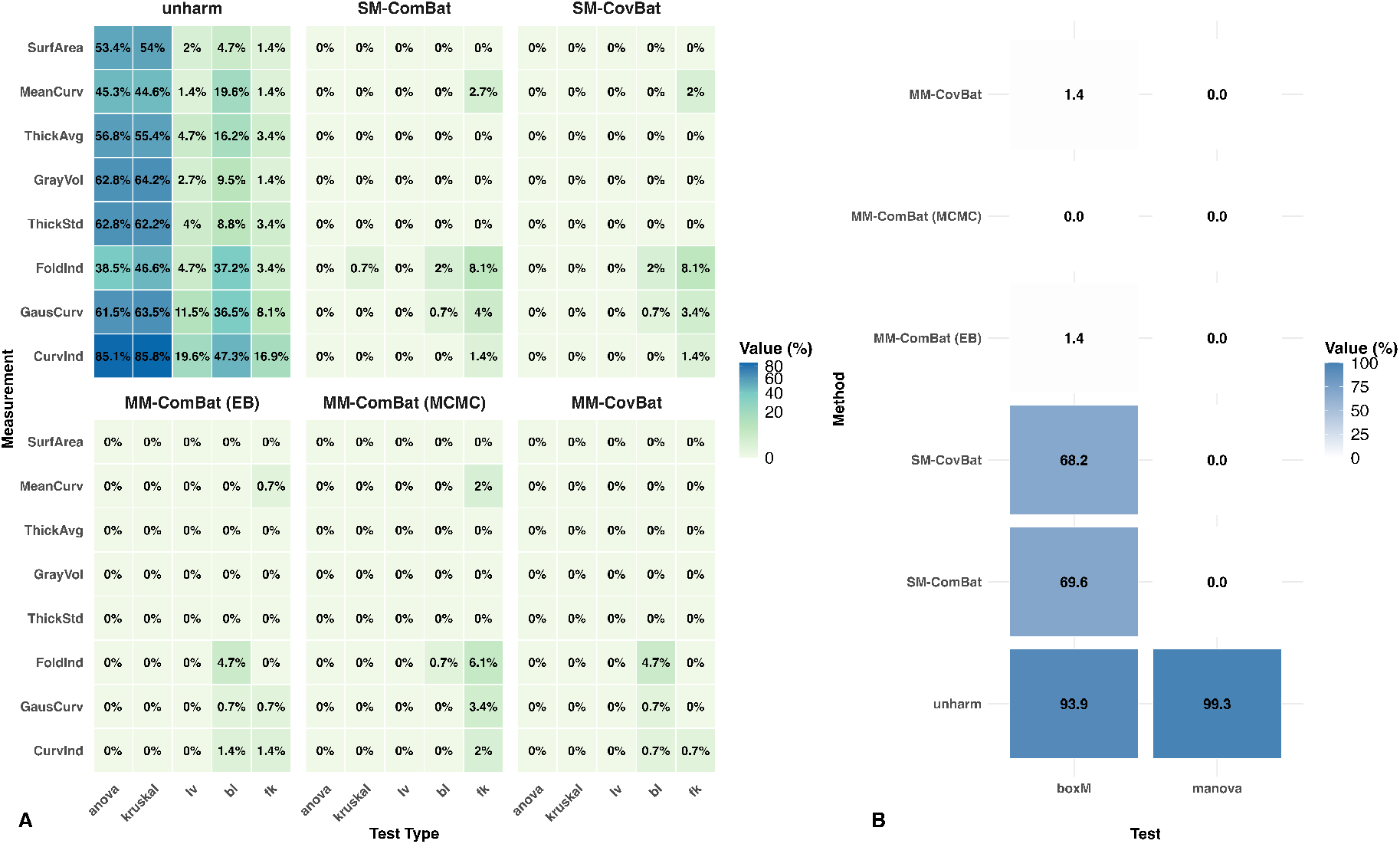
Statistical tests under the target-covariance formulation. (A) Univariate tests within each metric for additive and multiplicative batch effects across harmonization methods. (B) Multivariate tests of location and covariance across metrics. Overall, results are similar to those under the baseline covariance formulation, with slight improvement in within-metric scaling batch-effect removal for metrics with severe outliers.

**Figure S9.**
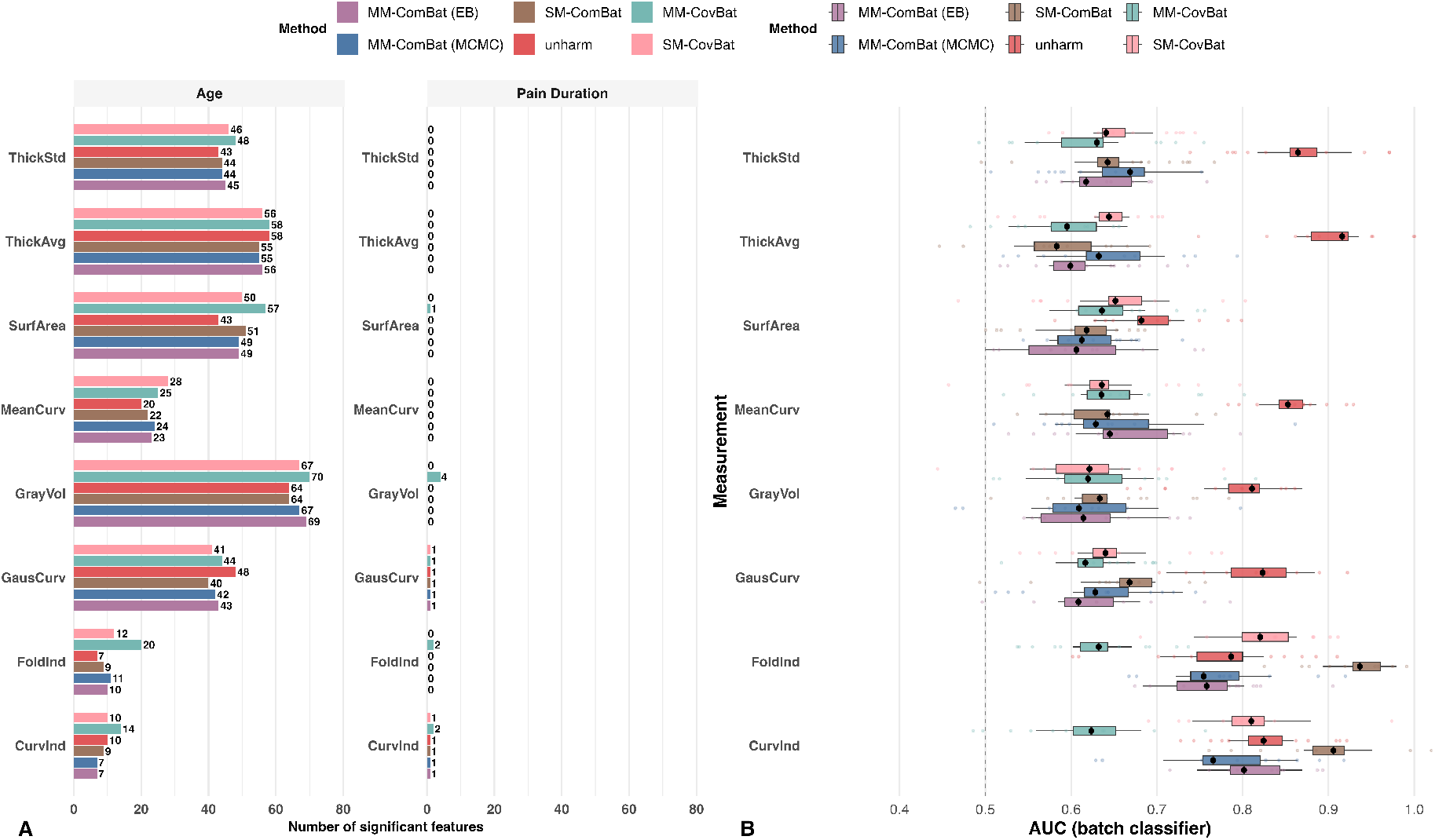
Detection of batch effects and biological variation under the target-covariance formulation. (A) Number of ROIs with significant biological associations after harmonization. (B) Residual global batch signal measured by 10-fold CV random-forest AUC for predicting site/scanner from harmonized features. Patterns are broadly consistent with those under the baseline covariance formulation, although apparent biological signal recovery is slightly attenuated and MM-ComBat retains some residual covariance-related batch signal for metrics with severe outliers.

**Figure S10.**
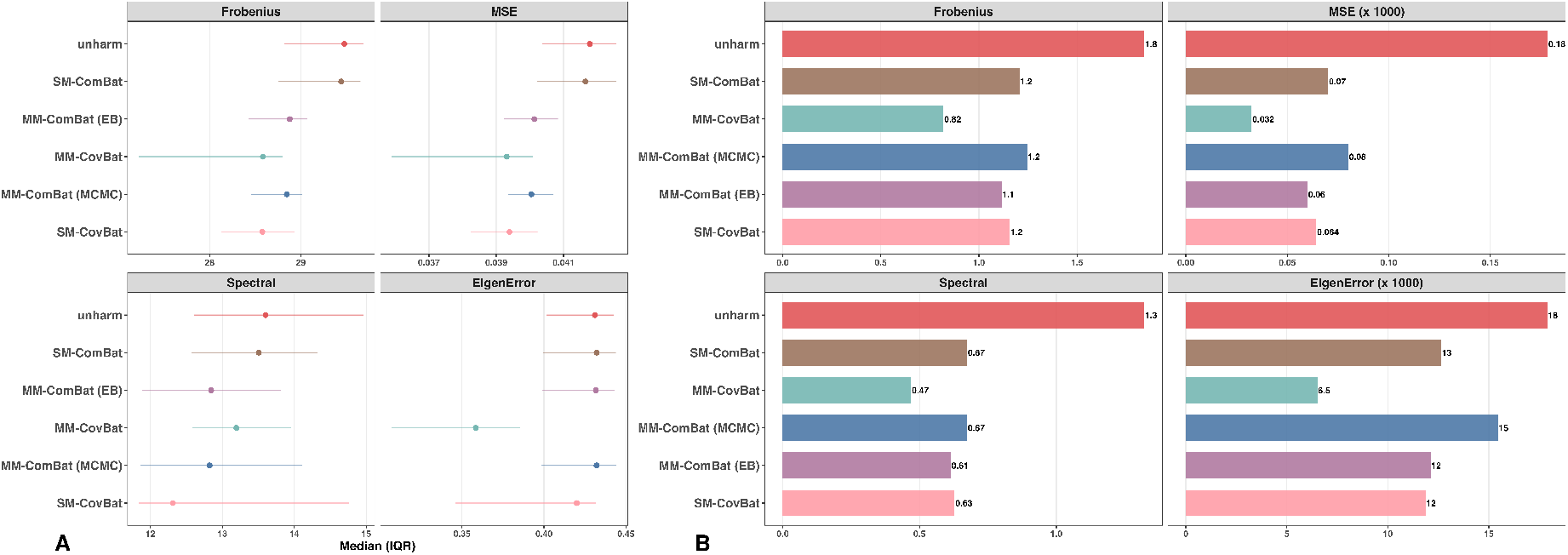
Batch detection in correlation under the target-covariance formulation. (A) Average within-metric correlation matrix distance across batch levels. (B) Average cross-metric correlation matrix distance across batch levels. The target-covariance formulation improves within-metric co-variance harmonization, especially for MM-CovBat, whereas performance for cross-metric covariance remains largely unchanged relative to the baseline covariance formulation.

**Figure S11.**
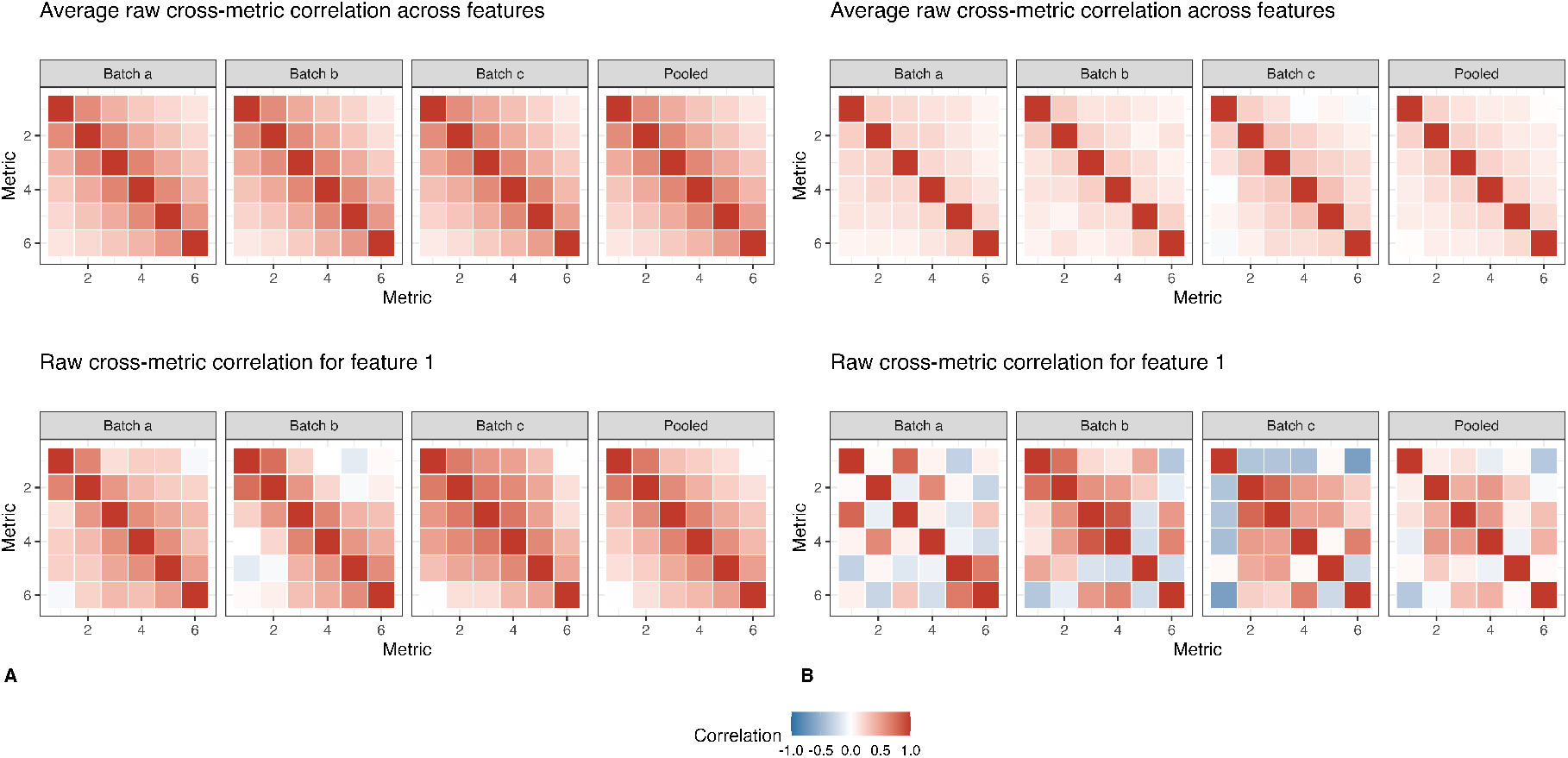
Diagnostic illustration of raw cross-metric correlation structure by batch. Panel (A) corresponds to a setting with strong biological structure, whereas panel (B) corresponds to a setting with strong batch structure. The top row shows the average raw cross-metric correlation matrix across features, and the bottom row shows the raw cross-metric correlation matrix for a representative feature (feature 1), each displayed separately by batch and in the pooled sample. Stable off-diagonal patterns in panel (A) reflect shared biological covariance across metrics, whereas the more diagonal average matrices and greater batch-specific heterogeneity in panel (B) indicate stronger batch-related distortion of cross-metric correlation structure.

### S4 Supplementary Table

**Table S1.**
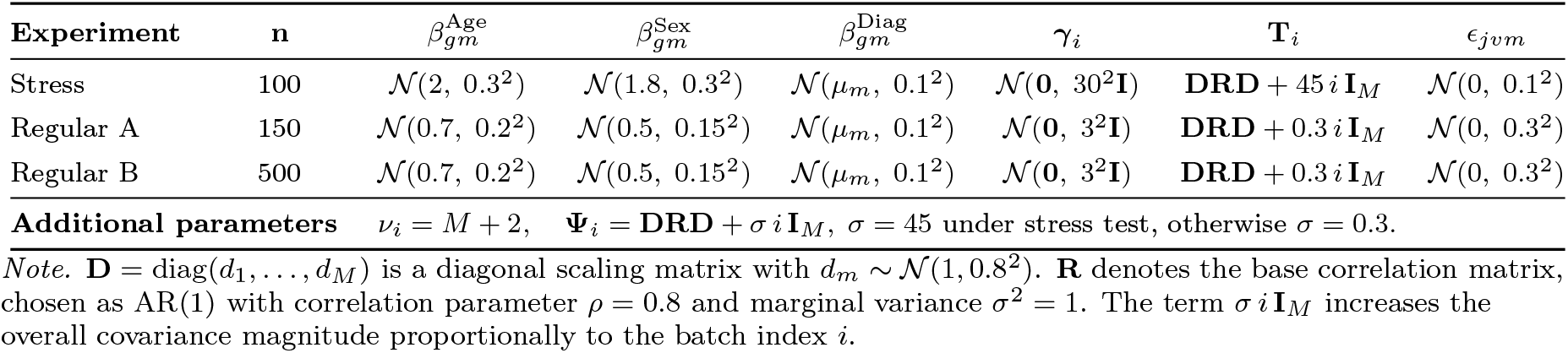
Three experimental conditions and parameter settings used across scenarios.

**Table S2.**
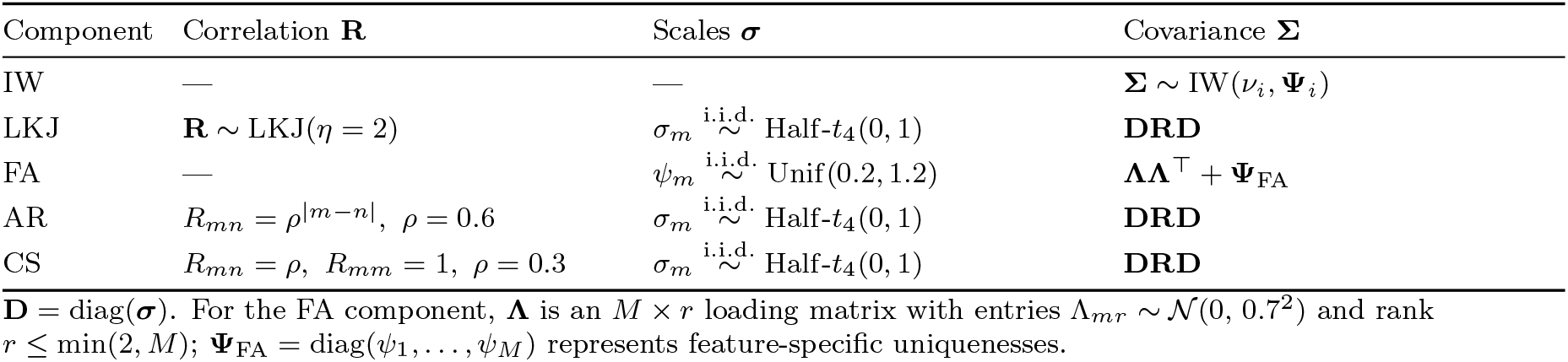
Covariance generators used in the model-misspecified mixture.

**Table S3.**
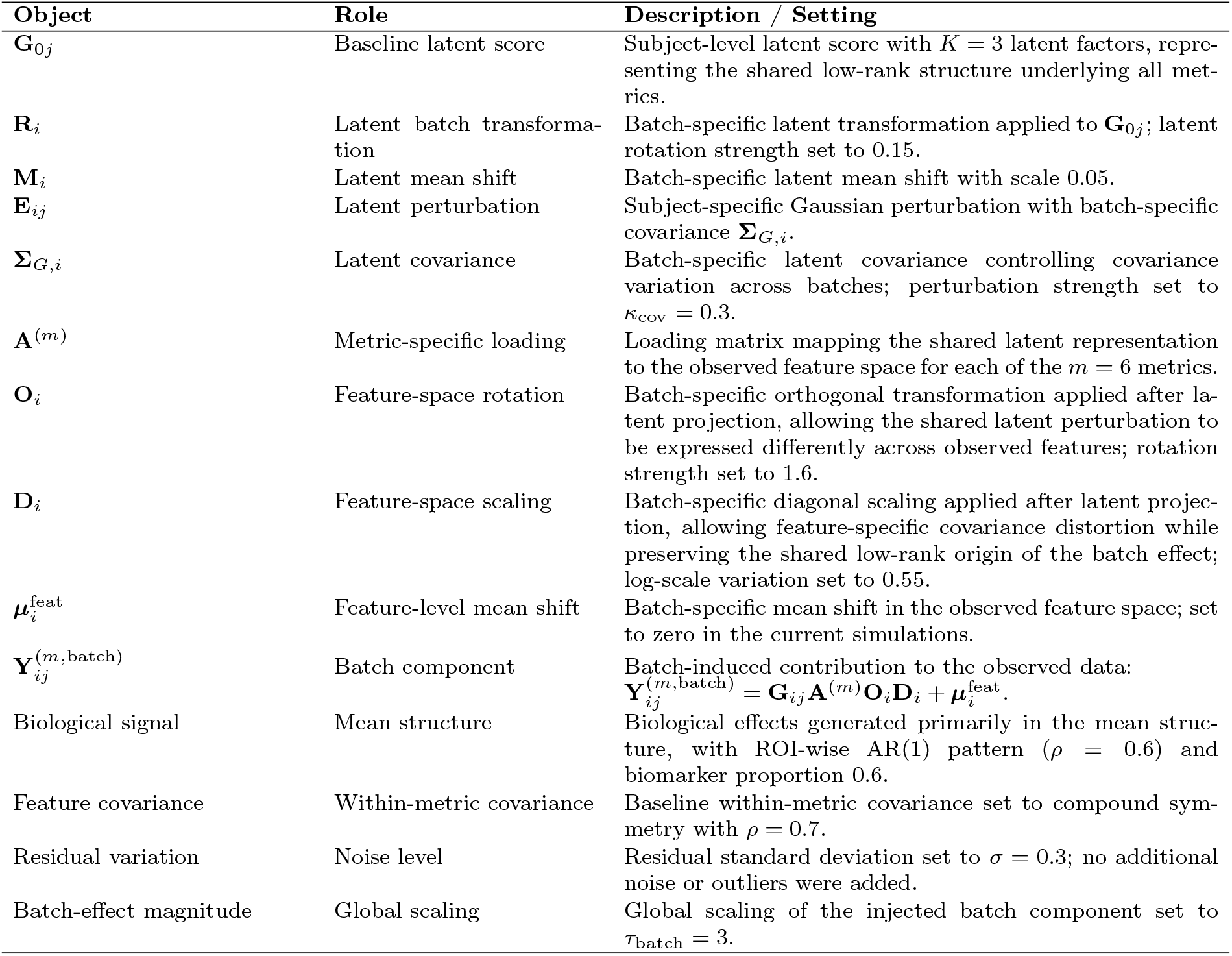
Core components of the latent-space batch-effect simulation.

**Table S4.** Simulation settings for biological covariance patterns across metrics.

| Scenario | $n$ | $\Sigma_v^{\text{bio}}$ | $\Delta_{iv}^{\text{batch}}$ | Mean shift | Noise | Outliers |
| --- | --- | --- | --- | --- | --- | --- |
| No batch / strong bio | 180 | AR(1), $\rho = 0.7$, $s = 1.5$ | 0 | 0 | 0.15 | None |
| Moderate batch / strong bio | 180 | AR(1), $\rho = 0.7$, $s = 1.7$ | Rotation, $s = 1.0$ | 1.0 | 0.20 | None |
| Moderate batch / strong bio + outliers | 180 | AR(1), $\rho = 0.7$, $s = 1.2$ | Rotation, $s = 1.0$ | 1.0 | 0.20 | 10%, outlier SD = 8 |
| Strong batch / weak bio | 180 | AR(1), $\rho = 0.7$, $s = 0.2$ | Low-rank, $s = 2.0$ | 1.5 | 0.20 | None |

*Note.* All scenarios use $K = 3$ batches, $M = 6$ metrics, and balanced batch assignment. Residual covariance is generated as $\Sigma_{iv}^{\text{res}} = \Sigma_v^{\text{bio}} + \Delta_{iv}^{\text{batch}} + \Sigma_{iv}^{\text{noise}}$ . Here, $s$ denotes the covariance-strength parameter. Biological effects are introduced in 30% of features. Rotation-based perturbations alter the orientation of the covariance structure across metrics, whereas low-rank perturbations introduce batch effects concentrated in a small number of dominant covariance directions.

**Table S5.**
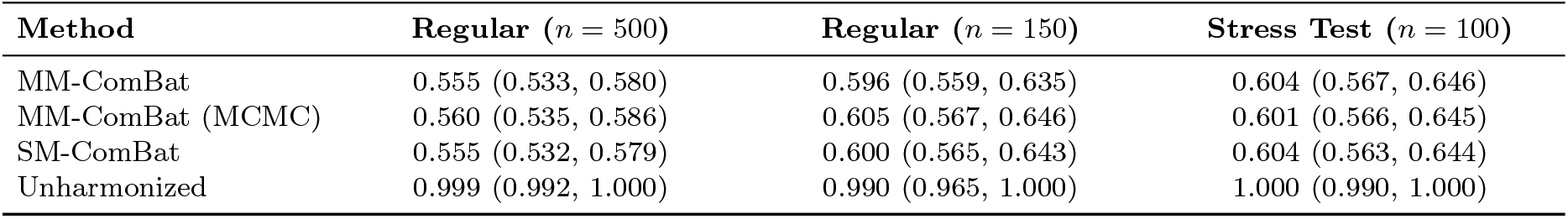
Within-metric batch-effect removal performance across experimental conditions (point estimates with 95% confidence intervals) under the model-misspecified scenario.

| Method | Regular ($n = 500$) | Regular ($n = 150$) | Stress Test ($n = 100$) |
| --- | --- | --- | --- |
| MM-ComBat | 0.555 (0.533, 0.580) | 0.596 (0.559, 0.635) | 0.604 (0.567, 0.646) |
| MM-ComBat (MCMC) | 0.560 (0.535, 0.586) | 0.605 (0.567, 0.646) | 0.601 (0.566, 0.645) |
| SM-ComBat | 0.555 (0.532, 0.579) | 0.600 (0.565, 0.643) | 0.604 (0.563, 0.644) |
| Unharmonized | 0.999 (0.992, 1.000) | 0.990 (0.965, 1.000) | 1.000 (0.990, 1.000) |

**Table S6.** Prior distributions and MCMC configuration for the multivariate Bayesian model.

| Component | Parameter | Description | Prior / Setting |
| --- | --- | --- | --- |
| Hyperparameters | $\nu_\sigma$ | Degrees of freedom for Half-$t$ prior on $\sigma_{iv}$ | Gamma(2, 0.1) |
| | $s_\sigma$ | Scale parameter for Half-$t$ prior | Cauchy(0, 2.5) |
| Batch-level effects | $\gamma_i$ | Batch-level intercept for batch $i$ | $\mathcal{N}(0, 1)$ |
| Mean structure | $\mu_{iv}$ | Feature- and batch-specific mean vector ($\in \mathbb{R}^M$) | $\mathcal{N}(\gamma_i \mathbf{1}, 2^2 \mathbf{I})$ |
| Scale parameters | $\sigma_{iv}$ | Feature- and batch-specific standard deviations ($\in \mathbb{R}_+^M$) | $t_{\nu_\sigma}^+(0, s_\sigma)$ |
| Correlation structure | $\mathbf{L}_{R,iv}$ | Cholesky factor of correlation matrix | LKJ_corr_chol(1) |
| Covariance construction | $\Sigma_{iv}$ | Feature- and batch-specific covariance matrix | $\Sigma_{iv} = \mathbf{L}_{\Sigma,iv} \mathbf{L}_{\Sigma,iv}^\top$ |
| | $\mathbf{L}_{\Sigma,iv}$ | Cholesky factor of covariance matrix | $\mathbf{L}_{\Sigma,iv} = \text{diag}(\sigma_{iv}) \mathbf{L}_{R,iv}$ |
| Likelihood | $\mathbf{Z}_{ijv}$ | Observed measurement vector for subject $j$ ($\in \mathbb{R}^M$) | $\mathcal{N}(\mu_{iv}, \Sigma_{iv})$ |
| MCMC sampler | — | Sampling algorithm | Hamiltonian Monte Carlo (Stan; NUTS) |
| Number of chains | — | Independent MCMC chains | 3 |
| Parallel chains | — | Chains run in parallel | 3 |
| Adaptation | — | Target acceptance probability | 0.98 |
| Diagnostics | — | Convergence assessment | $\hat{R}$ and effective sample size (ESS) |
| Posterior summaries | — | Reported estimates | Posterior means |

## References

Avants, Brian B. et al. (Feb. 2011). “A reproducible evaluation of ANTs similarity metric performance in brain image registration”. In: NeuroImage 54.3, pp. 2033–2044. issn: 1053-8119. doi: 10.1016/j.neuroimage.2010.09.025. url: http://dx.doi.org/10.1016/j.neuroimage.2010.09.025.

Barnard, John, Robert E. McCulloch, and Xiao-Li Meng (2000). “Modeling covariance matrices in terms of standard deviations and correlations, with application to shrinkage”. In: Statistica Sinica 10.4, pp. 1281–1311.

Bartlett, Maurice Stevenson (1937). “Properties of sufficiency and statistical tests”. In: Proceedings of the Royal Society of London. Series A-Mathematical and Physical Sciences 160.901, pp. 268–282.

Bayer, Johanna M. M. et al. (Oct. 2022). “Site effects how-to and when: An overview of retrospective techniques to accommodate site effects in multi-site neuroimaging analyses”. In: Frontiers in Neurology 13. issn: 1664-2295. doi: 10.3389/fneur.2022.923988. url: http://dx.doi.org/10.3389/fneur.2022.923988.

Beer, Joanne C. et al. (Oct. 2020). “Longitudinal ComBat: A method for harmonizing longitudinal multi-scanner imaging data”. In: NeuroImage 220, p. 117129. issn: 1053-8119. doi: 10.1016/j.neuroimage.2020.117129. url: http://dx.doi.org/10.1016/j.neuroimage.2020.117129.

Berardi, Giovanni et al. (Apr. 2022). “Multi-Site Observational Study to Assess Biomarkers for Susceptibility or Resilience to Chronic Pain: The Acute to Chronic Pain Signatures (A2CPS) Study Protocol”. In: Frontiers in Medicine 9. issn: 2296-858X. doi: 10.3389/fmed.2022.849214. url: http://dx.doi.org/10.3389/fmed.2022.849214.

Betancourt, Michael (2017). “A Conceptual Introduction to Hamiltonian Monte Carlo”. In: arXiv preprint 1701.02434. doi: 10.48550/ARXIV.1701.02434. url: https://arxiv.org/abs/1701.02434.

Box, G. E. P. (1953). “NON-NORMALITY AND TESTS ON VARIANCES”. In: Biometrika 40.3–4, pp. 318–335. issn: 1464-3510. doi: 10.1093/biomet/40.3-4.318. url: http://dx.doi.org/10.1093/biomet/40.3-4.318.

Carmon, Jona et al. (Oct. 2020). “Reliability and comparability of human brain structural covariance networks”. In: NeuroImage 220, p. 117104. issn: 1053-8119. doi: 10.1016/j.neuroimage.2020.117104. url: http://dx.doi.org/10.1016/j.neuroimage.2020.117104.

Chen, Andrew A., Joanne C. Beer, et al. (Dec. 2021). “Mitigating site effects in covariance for machine learning in neuroimaging data”. In: Human Brain Mapping 43.4, pp. 1179–1195. issn: 1097-0193. doi: 10.1002/hbm.25688. url: http://dx.doi.org/10.1002/hbm.25688.

Chen, Andrew A. and Matthew Gardner (2024). andy1764/ComBatFamily: v0.2.1. Version v0.2.1. doi: 10.5281/zenodo.13769617. url: https://github.com/andy1764/ComBatFamily.

Esteban, Oscar et al. (Dec. 2018). “fMRIPrep: a robust preprocessing pipeline for functional MRI”. In: Nature Methods 16.1, pp. 111–116. issn: 1548-7105. doi: 10.1038/s41592-018-0235-4. url: http://dx.doi.org/10.1038/s41592-018-0235-4.

Fischl, Bruce (Aug. 2012). “FreeSurfer”. In: NeuroImage 62.2, pp. 774–781. issn: 1053-8119. doi: 10.1016/j.neuroimage.2012.01.021. url: http://dx.doi.org/10.1016/j.neuroimage.2012.01.021.

Fligner, Michael A. and Timothy J. Killeen (Mar. 1976). “Distribution-Free Two-Sample Tests for Scale”. In: Journal of the American Statistical Association 71.353, pp. 210–213. issn: 1537-274X. doi: 10.1080/01621459.1976.10481517. url: http://dx.doi.org/10.1080/01621459.1976.10481517.

Fortin, Jean-Philippe, Nicholas Cullen, et al. (Feb. 2018). “Harmonization of cortical thickness measurements across scanners and sites”. In: NeuroImage 167, pp. 104–120. issn: 1053-8119. doi: 10.1016/j.neuroimage.2017.11.024. url: http://dx.doi.org/10.1016/j.neuroimage.2017.11.024.

Fortin, Jean-Philippe, Drew Parker, et al. (Nov. 2017). “Harmonization of multi-site diffusion tensor imaging data”. In: NeuroImage 161, pp. 149–170. issn: 1053-8119. doi: 10.1016/j.neuroimage.2017.08.047. url: http://dx.doi.org/10.1016/j.neuroimage.2017.08.047.

Gabry, Jonah et al. (2024). CmdStanR: The R Interface to CmdStan. url: https://mc-stan.org/cmdstanr/.

Hoffman, Matthew D. and Andrew Gelman (2014). “The No-U-Turn Sampler: Adaptively Setting Path Lengths in Hamiltonian Monte Carlo”. In: Journal of Machine Learning Research 15, pp. 1593–1623.

Horng, Hannah et al. (Mar. 2022). “Generalized ComBat harmonization methods for radiomic features with multi-modal distributions and multiple batch effects”. In: Scientific Reports 12.1. issn: 2045-2322. doi: 10.1038/s41598-022-08412-9. url: http://dx.doi.org/10.1038/s41598-022-08412-9.

Johnson, W. Evan, Cheng Li, and Ariel Rabinovic (Apr. 2006). “Adjusting batch effects in microarray expression data using empirical Bayes methods”. In: Biostatistics 8.1, pp. 118–127. issn: 1465-4644. doi: 10.1093/biostatistics/kxj037. url: http://dx.doi.org/10.1093/biostatistics/kxj037.

Kruskal, William H. and W. Allen Wallis (Dec. 1952). “Use of Ranks in One-Criterion Variance Analysis”. In: Journal of the American Statistical Association 47.260, pp. 583–621. issn: 1537-274X. doi: 10.1080/01621459.1952.10483441. url: http://dx.doi.org/10.1080/01621459.1952.10483441.

Levene, Howard et al. (1960). “Contributions to probability and statistics”. In: Essays in honor of Harold Hotelling 278, p. 292.

Lewandowski, Daniel, Dorota Kurowicka, and Harry Joe (Oct. 2009). “Generating random correlation matrices based on vines and extended onion method”. In: Journal of Multivariate Analysis 100.9, pp. 1989–2001. issn: 0047-259X. doi: 10.1016/j.jmva.2009.04.008. url: http://dx.doi.org/10.1016/j.jmva.2009.04.008.

Lu, Qianling et al. (Aug. 2024). “Shared and distinct cortical morphometric alterations in five neuropsychiatric symptoms of Parkinson’s disease”. In: Translational Psychiatry 14.1. issn: 2158-3188. doi: 10.1038/s41398-024-03070-z. url: http://dx.doi.org/10.1038/s41398-024-03070-z.

Orlhac, Fanny et al. (Jan. 2018). “A Postreconstruction Harmonization Method for Multicenter Radiomic Studies in PET”. In: Journal of Nuclear Medicine 59.8, pp. 1321–1328. issn: 2159-662X. doi: 10.2967/jnumed.117.199935. url: http://dx.doi.org/10.2967/jnumed.117.199935.

Pomponio, Raymond et al. (Mar. 2020). “Harmonization of large MRI datasets for the analysis of brain imaging patterns throughout the lifespan”. In: NeuroImage 208, p. 116450. issn: 1053-8119. doi: 10.1016/j.neuroimage.2019.116450. url: http://dx.doi.org/10.1016/j.neuroimage.2019.116450.

Reynolds, Maxwell et al. (2023). “ComBat Harmonization: Empirical Bayes versus fully Bayes approaches”. In: NeuroImage: Clinical 39, p. 103472. issn: 2213-1582. doi: 10.1016/j.nicl.2023.103472. url: http://dx.doi.org/10.1016/j.nicl.2023.103472.

Sadikov, Amir et al. (July 2025). “Mapping the microstructure of human cerebral cortex in vivo with diffusion MRI”. In: Communications Biology 8.1. issn: 2399-3642. doi: 10.1038/s42003-025-08523-9. url: http://dx.doi.org/10.1038/s42003-025-08523-9.

Sadil, Patrick et al. (2024). “Image Processing in the Acute to Chronic Pain Signatures (A2CPS) Project”. In: bioRxiv, pp. 2024–12.

Sluka, Kathleen A. et al. (June 2023). “Predicting chronic postsurgical pain: current evidence and a novel program to develop predictive biomarker signatures”. In: Pain 164.9, pp. 1912–1926. issn: 1872-6623. doi: 10.1097/j.pain.0000000000002938. url: http://dx.doi.org/10.1097/j.pain.0000000000002938.

Stan Development Team (2024). Stan User’s Guide. Version 2.38. url: https://mc-stan.org.

Storsve, A. B. et al. (June 2014). “Differential Longitudinal Changes in Cortical Thickness, Surface Area and Volume across the Adult Life Span: Regions of Accelerating and Decelerating Change”. In: Journal of Neuroscience 34.25, pp. 8488–8498. issn: 1529-2401. doi: 10.1523/jneurosci.0391-14.2014. url: http://dx.doi.org/10.1523/JNEUROSCI.0391-14.2014.

Tustison, Nicholas J et al. (June 2010). “N4ITK: Improved N3 Bias Correction”. In: IEEE Transactions on Medical Imaging 29.6, pp. 1310–1320. issn: 1558-254X. doi: 10.1109/tmi.2010.2046908. url: http://dx.doi.org/10.1109/TMI.2010.2046908.

Yu, Meichen et al. (July 2018). “Statistical harmonization corrects site effects in functional connectivity measurements from multi-site fMRI data”. In: Human Brain Mapping 39.11, pp. 4213–4227. issn: 1097-0193. doi: 10.1002/hbm.24241. url: http://dx.doi.org/10.1002/hbm.24241.

